# Catalytic inhibition of KAT6/KAT7 enhances the efficacy and overcomes primary and acquired resistance to Menin inhibitors in MLL leukaemia

**DOI:** 10.1101/2024.12.11.627663

**Authors:** Shellaina J. V. Gordon, Florian Perner, Laura MacPherson, Daniela V. Wenge, Wallace Bourgeois, Katie Fennell, Tabea Klaus, Jelena Petrovic, Jakub Horvath, Joan Cao, John Lapek, Sean Uryu, Jeffrey White, Enid Y. N. Lam, Xinmeng Jasmine Mu, Yih-Chih Chan, Andrea Gillespie, Benjamin Blyth, Michelle A. Camerino, Ylyva E. Bozikis, Henrietta Holze, Kathy Knezevic, Jesse Balic, Paul A. Stupple, Ian P. Street, Brendon J. Monahan, Shikhar Sharma, Elanor N. Wainwright, Dane Vassiliadis, Thomas A. Paul, Scott A. Armstrong, Mark A. Dawson

## Abstract

Understanding the molecular pathogenesis of MLL fusion oncoprotein (MLL-FP) leukaemia has spawned epigenetic therapies that have improved clinical outcomes in this often-incurable disease. Using genetic and pharmacological approaches, we define the individual and combined contribution of KAT6A, KAT6B and KAT7, in MLL-FP leukaemia. Whilst inhibition of KAT6A/B is efficacious in some pre-clinical models, simultaneous targeting of KAT7, with the novel inhibitor PF-9363, increases the therapeutic efficacy. KAT7 interacts with Menin and the MLL complex and is co-localised at chromatin to co-regulate the MLL-FP transcriptional program. Inhibition of KAT6/KAT7 provides an orthogonal route to targeting Menin to disable the transcriptional activity of MLL-FP. Consequently, combined inhibition rapidly evicts the MLL-FP from chromatin, potently represses oncogenic transcription and overcomes primary resistance to Menin inhibitors. Moreover, PF-9363 or genetic depletion of KAT7 can also overcome acquired genetic/non-genetic resistance to Menin inhibition. These data provide the molecular rationale for rapid clinical translation of combination therapy in MLL-FP leukaemia.

## INTRODUCTION

The mixed lineage leukaemia (MLL1; KMT2A) gene codes for a histone methyltransferase that plays an integral role in normal embryogenesis and the maintenance of both foetal and adult haematopoiesis(1–5). Recurrent chromosomal translocations involving this essential epigenetic regulator accounts for over 70% of infantile leukaemia and up to 10% of adult leukaemia(6). These MLL translocated leukaemias follow an aggressive clinical course with a poor response to conventional treatment, which has fuelled efforts to understand the molecular pathogenesis of the disease and use these insights to develop novel therapeutic strategies. MLL-fusion proteins (MLL-FP) are potent oncogenic drivers and provide one of the few examples in human cancer where a single genetic aberration is both necessary and sufficient for the full malignant phenotype. The importance of this fact means that even in the context of diverse subclonal genetic mutations, therapies that principally negate the oncogenic function of the MLL-FP can achieve meaningful clinical outcomes.

Central to the oncogenic function of MLL-FP is perturbed transcriptional regulation that manifests as the maintenance of a gene expression program that facilitates malignant self-renewal and impaired differentiation. A variety of approaches have been developed to curtail the transcriptional effects of MLL-FP including targeting the histone methyltransferase DOT1L which is aberrantly recruited by MLL-FP(7,8). Similarly, efforts to target transcriptional coactivators such as BET bromodomain family of proteins(9,10) or the YEATS domain proteins such as ENL(11,12) also showed substantial pre-clinical promise. Several of these targeted therapies proceeded to early phase clinical trials(13,14); however, for reasons that remain incompletely understood the clinical effects of these drugs as monotherapies were very modest.(15) More recently, small molecule inhibitors have been developed to interfere with the interaction between the amino-terminus of MLL1 and Menin, resulting in the displacement of the MLL-FP from the chromatin at key target genes and disrupting the gene expression program driven by this oncogene.(16,17). Importantly, the clinical evaluation of Menin inhibitors has shown significant promise even as single agents in heavily pre-treated patients with relapsed/refractory acute leukemias(18), validating the importance of these molecular insights and raising the possibility that Menin inhibition will form a key mainstay for future combination strategies in MLL-FP leukaemia.

Although therapeutic resistance ultimately emerged in patients treated with single agent Menin inhibitors(18,19), the clinical impact of this targeted therapy highlighted the importance of a detailed molecular rationale in guiding therapeutic intervention in this disease. In this regard, we and others recently demonstrated that in addition to binding Menin, the amino terminus of MLL also binds HBO1/KAT7(20,21). Moreover, targeting KAT7 either genetically and/or with proof-of-concept small molecule inhibitors had therapeutic potential in MLL-FP leukaemia(20,22). It has also been proposed that KAT6A, a related MYST acetyltransferase, through a functional interaction with ENL, may also play an important role in the molecular pathogenesis of MLL-FP leukaemia(23). In this study, we characterise a new selective catalytic inhibitor of KAT6A/B and KAT7. Using a variety of genetic and pharmacological approaches in a range of human and mouse MLL-FP leukaemia models we provide a detailed molecular rationale for targeting KAT6 and KAT7 together with Menin inhibition. We illustrate the importance of this combination to achieve a rapid and profound repression of the transcription program driven by MLL-FP leukaemia to achieve maximal therapeutic benefit.

## METHODS

### Chemicals

PF-9363 and PF-8144 was a gift from Pfizer. VTP50469 and SNDX-5613 (Revumenib) were purchased from MedChemExpress (cat. no. HY-114162, cat. no. HY-136175). For drug synergy characterisation experiments, cells were treated with DMSO (vehicle), 100 nM of VTP50469 and 2.5 µM of PF-9363 alone and in combination for 6 hr and 24 hr before taking samples for immunoblotting, RNA extraction or chromatin immunoprecipitation.

### Cell Culture

Molm-13 (RRID:CVCL_2119), THP-1 (RRID:CVCL_0006), K562 (RRID:CVCL_0004) and OCI-AML2 (RRID:CVCL_1619) cell lines were cultured in RPMI-1640 supplemented with 10% fetal bovine serum (FBS), 100 IU/ml Penicillin, 100 µg/ml streptomycin and 2 mM Glutamax. MV4;11 (RRID:CVCL_0064) cells were cultured in IMDM with 10% fetal bovine serum (FBS), 100 IU/ml Penicillin, 100 µg/ml streptomycin and 2 mM Glutamax. MLL-AF9 and MLL-AF9 LSC enriched cells were cultured in 20% fetal bovine serum (FBS), 100 IU/ml Penicillin, 100 µg/ml streptomycin, 2 mM Glutamax, mouse IL-3 (10 ng/ml) and mouse SCF (50 ng/ml) for the first week and then mouse IL-3 (10 ng/ml) for ongoing culture. Media for MLL-AF9 LSC was also supplemented with 1 µM iBET-151. MCF7 (RRID:CVCL_0031) cells were cultured in EMEM (ATCC, 30–2003) supplemented with 10% HI-FBS, 100 IU/ml Penicillin, 100 µg/ml streptomycin and 1X ITS-X at 0.01 mg/mL (Thermo Fisher #51500056). All cells were cultured in 5% CO2 at 37°C. All cell lines were regularly mycoplasma tested by PCR-analysis through the Peter MacCallum Cancer Centre Genotyping Core. Human cell lines were authenticated by short tandem repeat profiling through the Australian Genome Research Facility (Adelaide, South Australia).

### Immunoblotting

Cell nuclei were isolated in NEB buffer (15 mM Tris pH 7.5, 60 mM KCl, 15 mM NaCl, 5 mM MgCl2, 1 mM CaCl2, 250 mM sucrose supplemented with 10 mM sodium butyrate, 2 mM PMSF + 1X protease inhibitor cocktail, Roche). Histones were acid extracted in 0.2 M H2SO4 and precipitated in 33% trichloroacetic acid. 100 ng to 2 mg of purified histones were heated to 95°C in SDS sample buffer with 50 mM DTT for 5 min, separated by SDS-PAGE and transferred to 0.2 mm pore-size nitrocellulose membrane (Bio-Rad). The membranes were blocked in TBS-T buffer (Li-COR) and probed for indicated antibodies. Bands were visualised and analysed using the Li-COR CLX Odyssey machine and ImageStudioLite software.

### Antibodies

*Immunoblotting*: anti-mouse Total H3 (Abcam, ab10799, RRID:AB_470239, 1:5000), anti-mouse, Total H4 (Abcam, ab31830, RRID:AB_1209246, 1:1000), anti-rabbit H3K14ac (Abcam, ab52946, RRID:AB_880442, 1:1000), anti-rabbit H3K23ac (Merck, 07-355, RRID:AB_310546, 1:4000), anti-rabbit H4K12ac (Abcam, ab46983, RRID:AB_873859, 1:5000), anti-rabbit H4K5ac (Abcam, ab51997, RRID:AB_2264109, 1:100,000), goat anti-rabbit IRDye 680LT (Li-Cor, 926-68021, RRID:AB_10706309, 1:10,000), goat anti-mouse IRDye 800CW (Li-Cor, 926-32210, RRID:AB_621842, 1:10,000) *ChIP*: anti-rabbit Menin (Bethyl, A300-105A, RRID:AB_2143306), anti-rabbit MLL1(Bethyl, A300-086A, RRID:AB_242510), anti-rabbit IgG XP Isotype Control D1AE (CST, 3900S). 5 µg of antibody was used for MLL1, Menin and IgG isotype control immunoprecipitations.

*Cut&Tag:* anti-rabbit MLL1 antibody (Bethyl, A300-086A, RRID:AB_242510, 0.5µl/reaction), anti-rabbit KAT6A (Invitrogen, PA5-66742, RRID:AB_2664289, 0.5µl/reaction), anti-rabbit KAT7 (Abcam, ab190908, 0.5µl/reaction), anti-rabbit IgG control (Abcam, ab37415, RRID:AB_2631996, 0.5µl/reaction) and anti rabbit secondary antibody (Antibodies-Online ABIN101961, RRID:AB_10775589) *IP-MS:* anti-rabbit KAT7 antibody (Abcam, ab190908) and anti-rabbit IgG (R&D Systems, AB- 105-C, RRID:AB_354266)

*Flow Cytometry*: anti-mouse CD45.1, APC-Cy7 (Biolegend, clone A20, 110716, RRID:AB_313504), anti-mouse CD45.2, AF700 (Biolegend, Clone 104, 109822, RRID:AB_493730), anti-human HLA-A/B/C, AF647 (Biolegend, clone W6/32, 311416, RRID:AB_493135), Rat anti-mouse CD16/CD32 (BD, clone 2.4G2, 553141, RRID:AB_394656), anti-human CD11b antibody (Biolegend, Clone M1/70, 101212, RRID:AB_312794)

### Cell Proliferation Assay

Cells were seeded in triplicate in 12-well plates, and treated with either 25 nM PF-9363, 2.5 μM PF-9363 or DMSO (0.1%) over the indicated time. The drug was refreshed at every two days. Cells were stained with DAPI, and the live-cell number was calculated using the BD FACSVerse (BD Biosciences).

### AML MLL-FP Cell Proliferation assay

MLL-FP AML cell lines: EOL1 (RRID:CVCL_0258), OCI-AML2 (RRID:CVCL_1619), NOMO1 (RRID:CVCL_1609), ML2 (RRID:CVCL_1418), SHI1 (RRID:CVCL_2191), MV4;11 (RRID:CVCL_0064), Molm-14 (RRID:CVCL_7916), Molm-13 (RRID:CVCL_2119), OCI-AML4 (RRID:CVCL_5224) and THP-1 (RRID:CVCL_0006) were seeded in 24-well plate in complete cell culture media (1 mL/well) at optimized seeding densities for each cell line. The following day, three technical replicates were treated with a vehicle (DMSO) and a 10-point 3-fold serial dilution dose curve of PF-8144 with top dose of 3 µM. Every 3 to 4 days, additional 1 ml of fresh media with DMSO or corresponding PF-8144 drug concentration was added to the cells. At day 7, when vehicle-treated control wells reached near complete confluence, cells were split at appropriate ratio, according to the control well from each cell line and resuspended to make a total volume of 1 ml with replenished DMSO or PF-8144 in the same fashion as previously described. Cells were treated for a total of 21 days. After 21 days, cell viability was assessed using Cell Titer Glo Luminescent Cell Viability Assay (Promega) according to manufacturer’s instructions and plates were read using Envision Plate Reader. IC50 and AUC values were calculated using GraphPad Prism software.

### Drug Synergy Assay

10,000 cells were seeded into a 96-well plate. PF-9363 and VTP50469 were dispensed using the D300e Digital Dispenser (Tecan) and incubated for 96 hours. After 96 hours cells were resuspended in PBS and then split in half for Cell Titer Glo (Promega) assays according to manufacturer’s instructions or for propidium iodide staining to measure cell death. Cells were stained with propidium iodide (BD, 556463) at 1:400 before being analysed by flow cytometry on the BD LSR II. The percentage of PI positive cells in each sample was used as input for the synergyfinder R-package or synergyfinder web browser (https://synergyfinder.fimm.fi/) selecting the percent inhibition analysis option using default analysis options(24). Synergy scores were calculated using the Bliss model of drug synergy scoring(25). Each experiment was repeated in biological triplicate with each drug condition performed in technical duplicate on the plate.

### Myeloid differentiation synergy studies

OCI-AML2 (RRID:CVCL_1619) cells were seeded at 1.5 × 10^5^ in 10 cm cell culture dishes with 10 ml of complete cell culture media (RPMI supplemented with 10% heat-inactivated FBS and 100 IU/mL penicillin and 100 μg/mL streptomycin). The following day, cells were treated in 4 × 4 matrix of increasing concentrations of PF-8144 (6 nM, 20 nM and 60 nM) and SNDX-5613 (20 nM, 100 nM and 500 nM) with single compounds and their combinations for 7 days. At day 3, 10 ml of fresh media was added, and compounds were replenished (DMSO, PF-8144 and SNDX- 5613). After 7-days treatment cells were stained with anti-CD11b antibody (Clone M1/70, Biolegend, Cat. 101212, RRID:AB_312794) and DAPI before analysis by flow cytometry. CD11b frequency on live cells was used as input for synergyfinder to assess Bliss synergy scores(24,25).

### Quantitative real-time Polymerase Chain Reaction

RNA was extracted from drug treated cells using the Qiagen RNAeasy Kit. On column DNAse treatment was performed. cDNA was generated using the SuperScript Vilo cDNA synthesis kit (Life Technologies) according to manufacturer’s instructions. Quantitative PCR was performed using primers for *MEIS1* and *HOXA9* (Supplementary Table 1) on the StepOne Real-Time PCR system (Applied Biosystems) or Bio-Rad CFX qPCR system with SYBR green reagents. All samples were assayed in triplicate. Expression levels were determined using the ΔΔCt method(26) and normalised to *B2M*.

### SPLINTR barcoding

Lentivirus containing SPLINTR-V2-EF1a_GFP barcode library was produced as previously described(27). Molm-13 (RRID:CVCL_2119) cells were transduced with SPLINTR-V2- EF1a_GFP barcode library at a low multiplicity of infection, generating a population of 5-10% GFP-positive cells. After 48 hours, barcode-positive cells were sorted using the BD Fusion 5 to isolate a pure population. 15K cells were plated and expanded for 2 weeks or ∼12 doublings to ensure equal representation of barcodes for all mice in the *in vivo* experiment. Each mouse received 1 × 10^6^ barcoded cells via tail vein injection. 1 × 10^5^ cells were lysed in Viagen lysis buffer (Viagen Biotech) with 0.5 mg/ml proteinase K (Invitrogen) according to manufacturer’s instructions for DNA-barcode sequencing. The remaining cells were cryopreserved in 10% DMSO in FBS as a baseline (T0) sample for scRNA-sequencing.

### Mouse Details

All animal experiments were conducted according to regulatory standards approved by the Peter MacCallum Cancer Centre Animal Ethics and Experimentation committee or at the Dana-Farber Cancer Institute under the oversight of the Institutional Animal Care and Use Committee (IACUC) protocol #16-021. All procedures performed on animals were in accordance with regulations and established guidelines and were reviewed and approved by an Institutional Animal Care and Use Committee.

### Mouse Experiments

1 × 10^5^ mouse mCherry positive MLL-AF9 cells were injected into the tail vein of NOD.Cg- *Prkdc^scid^ Il2rg^tm1Wjl^*/SzJ (NSG, RRID:IMSR_JAX:005557) mice aged 6-10 weeks. Ten days following transplantation, mice were randomised into treatment groups (n=6) based on leukaemia burden (mCherry-positive cells) detected by flow cytometry in the peripheral blood and monitored weekly thereafter. Mice were dosed once daily with Vehicle (5% DMSO/40% PEG300/55% PBS), 0.3 mg/kg PF-9363 or 5 mg/kg PF-9363 via oral gavage. Mice reached endpoint upon displaying altered gait or leukaemia burden >10%. For the Molm-13 xenograft experiment, 1 × 10^6^ SPLINTR barcoded Molm-13 cells were injected into the tail vein of NSG mice aged 6-10 weeks. Seven days following transplantation, mice were randomised into treatment groups (n=5) based on leukaemia burden measured by flow cytometry (HLA-A/B/C-positive cells). Mice were dosed twice daily with 10 mg/kg of SNDX-5613, once daily with 5 mg/kg PF-9363 or vehicle via oral gavage for about 21 days (42 oral gavage ethical limit). Peripheral blood leukaemia burden was monitored weekly. Mice were culled after displaying altered gait. At endpoint, 5 × 10^5^ cells from the bone marrow and spleen of each mouse were harvested for SPLINTR barcode sequencing and additional cells were cryopreserved for scRNA-seq.

### Patient Derived Xenograft Model

Patient-derived xenograft (PDX) models of leukaemia were previously established, characterized and banked at Dana-Farber Cancer Institute’s Center for Pediatric Cancer Therapeutics (CPCT). Leukaemia cells cryopreserved from each graft were injected into non-conditioned NOD.Cg- Prkdc^scid^Il2rg^tm1Wjl^/SzJ (NOG, RRID:IMSR_TAC:HSCFTL-NOG) aged 8-12 weeks at a dose of 2.5 × 10^5^ cells per mouse. Engraftment progress was monitored in the peripheral blood using flow cytometry every 3-4 weeks, detecting the ratio of human CD45 (labelled with PE) to mouse CD45 (labelled with APC-Cy7) on a LSRFortessa instrument (BD Biosciences). Upon detection of engraftment oral administration of 0.033% or 0.1% VTP50469 (supplemented rodent special diet) or 5 mg/kg PF-9363 (once daily, intraperitoneal) was initiated. Mice were continuously monitored for peripheral blood chimerism of human cells. Mice that reached the predefined study endpoint were humanely euthanized using CO2 inhalation followed by cervical dislocation. Bone marrow cells were extracted from cleaned tibias, femurs, iliac crests, and lumbar vertebrae. Spleen cells were obtained by passing the organ through a 40 μM nylon cell filter (FALCON). To measure the burden of human leukaemia cells in blood, spleen, and bone marrow, flow cytometry was utilized, quantifying the ratio of human CD45 (labelled with PE) to mouse CD45 (labelled with APC-Cy7) all of which were supplied by Biolegend.

### CRISPR-Cas9 cell competition assay

Cell lines were lentivirally transduced using Lenti-Cas9-2A-blast (RRID:Addgene_73310) to achieve stable expression of *Streptococcus pyogenes* Cas9. Following transduction, the cells were selected using Blasticidin (5-10 µg/ml). Subsequently, lentiviral particles containing specific guide RNAs in the pUSEPR vector system, were introduced to the cells. Target cells were exposed to the RFP-expressing vector at a moderate to low Multiplicity of Infection (MOI), leading to the emergence of a population with 20-50% RFP-positive (guide-expressing) cells. For the cell competition experiments, target cells were cultured in 96-well plates. The fraction of RFP-positive cells, which represented the population expressing the guide RNAs, was initially assessed at baseline (3 days after transduction) and at the specified time points using a BD LSR Fortessa flow cytometer. The ratios between the baseline measurement and the corresponding time points were computed, and these ratios were then presented as relative fractions (compared to the baseline chimerism).

### SPLINTR Barcode-seq and Analysis

At disease endpoint, 5 × 10^5^ cells from bone marrow and spleen of NSG mice were lysed in Viagen lysis buffer (Viagen Biotech) with 0.5 mg/ml proteinase K (Invitrogen) according to manufacturer’s instructions. Barcode sequences were amplified using primers flanking the constant region of the barcode before adding i5 and i7 indexes compatible with NGS sequencing as described previously^3^. Samples were sequenced on an Illumina NextSeq2000 using 100-bp single end chemistry targeting 2 million reads per sample. Fastq files were processed using the BARtab v1.4 pipeline (https://github.com/DaneVass/BARtab) for bulk population analysis as previously described(28). The bartools v1.0 R package (https://github.com/DaneVass/bartools) was used for further downstream quality control using default settings before analysis and visualisation(28). For differential barcode abundance calculations, PCR replicates were averaged and collapsed before using the barbie package to filter for barcodes in the top 90^th^ percentile of any mouse sample. The voom transformation(29) was applied to the filtered barcode counts data. Limma (30) (RRID:SCR_010943) was used to identify statistically significant (p-value < 0.05 and absolute log2 fold change of 1) differences in SPLINTR barcode abundance between combination and vehicle treatment groups. The results of the differential barcode abundance analysis were visualised using EnhancedVolcano (31) in R.

### RNA-seq and Analysis

RNA was extracted from drug treated cells using the Qiagen RNAeasy Kit. RNA concentration was determined using Qubit RNA High Sensitivity kit (Thermo Fisher Scientific). Libraries for the synergy characterisation experiment was prepared using the QuantSeq 3’ mRNA-seq Library Prep kit (Lexogen) and sequenced on the NextSeq2000 system using 100-bp single end chemistry targeting 10 million reads per sample. Libraries for genetic deletion of MYST KATs in mouse MLL-AF9 cells were prepared using the NEBNext-UtraII mRNAseq Library Prep and sequenced on the NextSeq500 system using 37bp paired end chemistry targeting 10 million reads per sample. Fastq files were generated using Bcl2fastq (v2.20.0.422, Illumina, RRID:SCR_015058). Adapters were trimmed from raw reads using Trim Galore v0.4.3 (RRID:SCR_011847). Trimmed reads were aligned to the human genome (GRCh38.78) using HiSAT2 v2.2.1 (RRID:SCR_015530) and assigned to genes using htseq-count (RRID:SCR_011867) (32,33). Genes were annotated using the biomaRt R package (v2.48.3, RRID:SCR_019214) and differential expression was calculated using DESeq2 v1.40.2 (RRID:SCR_015687) (34). Genes with a false-discovery rate, corrected for multiple method of Benjamini and Hochberg, below 0.05 and an absolute log2 transformed fold change of 1 or greater were considered significantly differentially expressed. Volcano plots were generated using ggplot2, heatmaps were generated using pheatmap (RRID:SCR_016418), and gene set enrichment plots were generated using fgsea v1.28.0 (RRID:SCR_020938) (35).

For analysis pertaining to genetic knockouts in the MLL-AF9 cells, genes with an adjusted p-value < 0.05 and absolute log2 fold change of 1.5 were identified as statistically significant. Clusters of differentially expressed genes between samples were identified using K-means clustering where K = 6.

### Cut and Tag

The protein A–Tn5 fusion protein (pA–Tn5) was produced using the 3XFlag-pA-Tn5-Fl plasmid (Addgene, 124601) in a bacterial expression system and Cut&Tag was performed as previously described(36). In short, 1 × 10^5^ cells were washed with PBS and then with a wash buffer (20 mM HEPES, pH 8.0, 150 mM NaCl, 0.5 mM spermidine), followed by immobilization on concanavalin A (ConA) magnetic beads activated in binding buffer (20 mM HEPES, pH 8.0, 10 mM KCl, 1 mM CaCl2, 1 mM MnCl2). Bead-bound cells were then permeabilized by washing with wash buffer supplemented with NP40 (0.01%) and Digitonin (0.01%) and incubated with anti-MLL1 antibody (Bethyl, A300-086A), anti-KAT6A (Invitrogen, PA5-66742), anti KAT7 (Abcam, ab190908) or rabbit IgG control (Abcam, ab37415) at 4°C. After a 2 hr incubation, the cells were washed twice and incubated with a secondary antibody (Antibodies-Online ABIN101961) for 1 hr at room temperature (RT). After 2 additional washes, cells were loaded with the pA–Tn5 conjugate in wash buffer supplemented with NP40 (0.01%), Digitonin (0.01%) and 300mM NaCl and incubated for 1 hr at RT. Cell-bead complexes were then washed 3 times and resuspended in 50 µl of tagmentation buffer (20 mM HEPES, pH 8.0, 300 mM NaCl, 0.5 mM spermidine, 0.01% Digitonin, 0.01% NP40, 10 mM MgCl2) at 37°C for 1 hr to allow tagmentation of antibody-labelled regions. Subsequently, tagmentation was stopped by addition of STOP buffer (10 mM Tris, 40 mM EDTA, 0.2% SDS, 0.4 mg/ml Proteinase K) and incubated at 37°C overnight. DNA was extracted by purification with AmpureXP (1:1) and libraries were constructed as previously described(36).

### Immunoprecipitation Mass Spectrometry

MCF7 cells were seeded in triplicates to a 100mm tissue culture dish and grown in corresponding growth media until confluency of about 1 × 10^7^ cells. Cells were washed and collected by scraping before proceeding to nuclear extraction using Active Motif Nuclear Complex Co-IP kit (Cat. No 54001) following manufacturer’s protocol. Briefly, cells were resuspended in pre-chilled 500µl 1x Hypotonic Buffer (supplemented with 2 mM Sodium Butyrate, 1 mM DTT) and incubated on ice for 15 min. Next, 25µl of detergent was added and nuclei were pelleted. The supernatant was discarded, and nuclei pellet was resuspended in 100µl of complete digestion buffer with 0.5µl enzymatic shearing cocktail and incubated at 4°C for 90 min. The reaction was stopped with 2µl 0.5 M EDTA. Next, nuclear extracts were cleared at max speed for 10 min at 4°C. All of supernatant containing digested nuclear extract was transferred to a new tube and 500µl of Low IP buffer without additional NaCl or detergent was added together with 5-10µg of KAT7 antibody (Abcam #190908) or rabbit IgG (R&D Systems, AB-105-C) and incubated overnight at 4°C. The next day 50µl of prewashed Dynabeads Protein G beads were added to the mixture and incubated for an additional 2 hrs. Next, the samples were washed with Low IP buffer and post-wash the beads directly were subjected to downstream processing for Mass Spectrometry analysis. IP pulldowns were in parallel confirmed using western blotting.

Mass spectrometry was performed using a Thermo Orbitrap Fusion Lumos mass spectrometer. 4µl of each sample was injected onto a 75 µm × 50 cm, 2 µm C18 column (Thermo Scientific ES803A) and separated over a 105-minute gradient on an Easy nLC 1200 operated at 300 nL/min. The gradient was from 10% to 34% buffer B (80% Acetonitrile with 0.1% formic acid) for 105 minutes followed by a linear ramp to 100% buffer B in 5 minutes. After 10 minutes, the column was returned to initial conditions and equilibrated.

The mass spectrometer was operated in data dependent mode with a five second cycle. MS1 scans were acquired in the Orbitrap with a resolution of 60,000 over a range of 500-1200 m/z. Automatic gain control (AGC) target was set to 2 × 10^5^ with a maximum inject time of 100 ms. Peptides were chosen for fragmentation if they had a charge of 2-6 and an intensity of at least 5 × 10^4^. Dynamic exclusion was enabled for 90 seconds. All MS2 spectra were acquired in the linear ion trap, with the quadrupole used for isolation with a window of 0.5 m/z. The AGC target for fragmentation spectra was 1 × 10^4^ with rapid scan rate. The maximum injection time was 35 ms. Peptides were fragmented with CID at 30% normalized collision energy with an activation time of 10 ms and an activation Q of 0.25. For MS3 spectra, up to 10 ions were selected for synchronous precursor selection, and data were collected at 60,000 resolution in the Orbitrap. Ions were fragmented with HCD at an energy of 55%. MS3 AGC was set to 1 × 10^5^ with a maximum injection time of 250 ms and a first mass of 110 m/z. Data at all stages were centroided. Resultant raw files were processed on an IP2GPU server (Integrated Proteomics Applications, Inc.). Data were searched with the ProLuCID algorithm(37) against the Uniprot Human Database (Downloaded January 29, 2018) concatenated with the current contaminants database and reverse database. Carbamidomethylation of Cysteine residues (+57.02146) and TMTpro modification of peptide n-termini and Lysine residues (+304.2071) were included as static modifications. Oxidation of Methionine (+15.9949) was included as a variable modification. A maximum of 2 variable modifications and two missed cleavages were allowed. Peptides had to have a minimum length of 6 amino acids to be considered. Data were searched with a 50 ppm MS1 tolerance(38) and 600 ppm MS2 tolerance. Final data were filtered to a 1% protein level false discovery rate.

### Chromatin Immunoprecipitation

30 million cells were crosslinked with 2 mM disuccinimidyl glutarate (DSG, Sigma-Aldrich) for 30 minutes at room temperature followed by crosslinking with 1% v/v methanol-free formaldehyde (Pierce) for 15 minutes. Crosslinking was quenched by adding Tris HCl pH 8.0 to sample reaching a final concentration of 250 mM. Samples were washed twice with cold PBS before resuspending in ChIP shearing buffer (50 mM Tris HCl pH 8.0, 1% SDS, 10 mM EDTA with protease and phosphatase inhibitors, Roche). Samples were sonicated using a Covaris ME220 for 7.5 minutes using the following sonication conditions: PIP 75, CPB 1000, Duty Factor 15%, temperature 7°C (min/max 6/12). Samples were centrifuged at 10°C for 15 minutes at 20,000 g before quantifying the clarified supernatant using the Qubit dsDNA Broad Range kit (Thermo Fisher Scientific). 5% total chromatin was used as input and 30 µg of chromatin was used for Menin and MLL1 immunoprecipitations. Chromatin was diluted 1:10 with wash buffer (50 mM Tris-HCl pH 8.0, 166 mM NaCl, 1.11% Triton-X 100, 2 mM EDTA) and incubated with 5 µg MLL1 or Menin antibodies overnight at 4°C on a rotator. Antibody bound chromatin was incubated with Protein A beads (Thermo Fisher) for 4 hours before washing. Antibody bound beads were placed on a magnetic rack and washed 5 times with wash buffer, once with high salt wash buffer (50 mM Tris-HCl pH 8.0, 500 mM NaCl, 1.11% Triton-X 100, 2 mM EDTA) and once with TE buffer (10 mM Tris-HCl pH 8.0 and 1 mM EDTA). Samples were resuspended in elution buffer (20 mM Tris-HCl pH 8.0, 5 mM EDTA, 200 mM NaCl, 1% SDS and 200 µg /ml Proteinase K) and incubated overnight at 65°C to reverse crosslinks. Immunoprecipitated DNA was purified using the Qiagen MiniElute PCR purification kit before quantification using the Qubit dsDNA High Sensitivity assay (Thermo Fisher Scientific). Enrichment of DNA encoding known MLL1 and Menin binding sites were validated using ChIP-qPCR prior to library preparation using the Takara DNA-Thruplex Library Prep kit. Samples were sequenced on the NextSeq 2000 P2 using 100-bp single end chemistry targeting 20 million reads per sample.

### ChIP-seq Analysis

Quality of reads were assessed with FastQC v0.11.6 (RRID:SCR_014583) and adapters were trimmed with Cutadapt (RRID:SCR_011841) (39) as implemented in Trim Galore v0.4.4 (RRID:SCR_011847). Reads were then aligned to the Hg38 reference genome using Bowtie2 v2.3.4.1 (RRID:SCR_016368) (40) with the --no-unal, --no-discordant and --no-mixed flags plus -I 10 and -X 700 in addition to default settings. Duplicates were removed by implementing the Picard toolkit v2.6.0. Sorting and indexing were performed with SAMtools v1.9 (RRID:SCR_002105) (41). Broad peaks were called with MACS2 v2.1.1 (42). RPGC normalised bigwigs and heatmaps were generated with deeptools v3.5.0 (RRID:SCR_016366) (43).

### Single Cell RNA-seq

scRNA-seq analysis of all samples was conducted using the 10X Genomics 3’ V3 Single Cell Gene Expression Library Preparation kit on a 10X Genomics chromium instrument. Cryopreserved SPLINTR barcoded cells from bone marrow and spleen of mice in the vehicle and combination treatment groups and baseline samples were thawed rapidly at 37 °C. Viable, propidium iodide negative and SPLINTR barcoded, GFP positive, cells were sorted using the BD Fusion 5. Cells were labelled with lipid-modified oligonucleotides (LMO) as previously described(44). Samples were pooled and resuspended so that the final cell concentration was suitable for loading on to the 10X Chromium Single Cell Chip (1,500 cells/µl). Two captures were performed and scRNAseq library generation was continued as per the maufacturers recommendations. Libraries were sequenced on the Illumina NovaSeq using paired-end 100-bp chemistry targeting 70,000 reads per cell. LMO labelled libraries were sequenced on the NextSeq2000 using single-end 100-bp chemistry targeting ∼5,000 reads per cell.

### Single Cell RNA-seq analysis

Cell x gene count matrices from each of the two captures were generated using the 10x Genomics Cell Ranger (v.7.1.0, RRID:SCR_017344) count pipeline against the Cell Ranger mm10/GRCm38 v3.0.0 genome(45). scRNA-seq quality control was performed using Seurat v5.1.0 (RRID:SCR_016341) (46) in R v4.3.2. Low-quality cells, defined as those with fewer than 200 genes or 4,000 unique molecular identifiers (UMIs) detected, were filtered out. Cells with greater than 20% mitochondrial RNA content were also removed. Multiplexed scRNA-seq samples containing TotalSeq-A hashtag oligo information were demultiplexed using the CITE-seq-count v.1.4.5 (RRID:SCR_019239) (47) and HTODemux methods within Seurat v.5.1.0 using default settings. For SPLINTR barcode detection from single-cell data, unmapped reads were extracted from BAM alignment files for each scRNA-seq library using SAMtools v.1.9 (RRID:SCR_002105) (41). BAM entries containing SPLINTR barcode reads were identified and extracted using BARtab v1.4(28). Sequenced PCR amplicons containing SPLINTR barcode information were also processed by BARtab using the single-cell workflow with fast input and cells called as non-empty droplets by Cell Ranger as whitelist. This resulted in a data frame of cell ID and SPLINTR barcode ID, which was loaded into corresponding Seurat scRNA-seq objects using the AddMetaData function. Where available, 10x scRNA-seq droplets containing doublet cells were identified based on hashtag oligo combinations with the HTODemux function in Seurat v5.1.0 and were removed. Hashtag barcodes corresponding to individual samples are provided in Supplementary Table 3. Cell cycle phase assignments were generated per cell from scores derived using the CellCycleScoring function within Seurat v5.1.0. A list of the genes used to determine cell cycle, and all other gene sets used in the scRNA-seq analysis is provided in Supplementary Table 4.

Filtered scRNA-seq datasets underwent log normalisation with the percentage of mitochondrial reads, S phase cell cycle score and G2M cell cycle phase score variables included in the regression model as sources of technical variation to remove except where indicated. Dimensional reduction, k-nearest neighbour graph construction and clustering were performed using Seurat v5.1.0. In general, the default settings were used unless specified otherwise. The first 50 principal components were used to compute a nonlinear dimensional reduction using the UMAP method(48). Louvain clustering was performed for a resolution range of 0.1–2 (0.1 step size). Marker gene identification and differential gene expression analysis for clusters and groups of cells was performed using Wilcoxon rank sum tests within Seurat v5.1.0. Gene set enrichment analysis was performed with fgsea v1.28.0 (RRID:SCR_020938) (35).

### Statistics and Reproducibility

Statistical analysis (t-test, one-way ANOVA) was carried out using GraphPad Prism v10. Details of the statistical analyses and significance values are specified in the figures and figure legends. Non-significant data is not annotated or denoted as ns. Data were reported as the mean ± SEM or independent replicates shown as individual data points.

## RESULTS

### Inhibition of KAT7 is required for the best therapeutic response

Although proof-of-concept small molecules that specifically target the MYST acetyltransferases KAT6A/B and KAT7 have been developed(20,49), these compounds were limited by their pharmacokinetic properties for use in pre-clinical mouse models. Consequently, whilst the genetic evidence for targeting KAT6A(23) or KAT7(20,22) in MLL-FP is compelling, there is limited evidence to support the catalytic inhibition of these targets *in vivo*. Moreover, in pre-clinical mouse models it is not known if catalytic inhibition of KAT6A/B is sufficient for therapeutic response or if concurrent inhibition of KAT7 provides greater efficacy. The pharmacokinetic limitations that have hindered progress in these studies have recently been overcome with a new chemical series(50), which potently inhibits KAT6A/B at low concentrations and KAT7 at higher concentrations (**Fig. 1A**). Although the substrate preference for the MYST acetyltransferases is promiscuous *in vitro*, it is now well established that in a cellular context H3K23ac is the major substrate for KAT6A/B(50) and H3K14ac is almost exclusively deposited by KAT7(20,51). Consistent with this we found that CTX-648/PF-9363 (hereafter referred to as PF-9363) selectively inhibited KAT6A/B at 25 nM in cells grown *in vitro* but a higher dose of 2.5 µM also inhibited KAT7 (**Fig. 1B**). In line with the rapid turnover of histone acetylation, we found that catalytic inhibition resulted in the rapid loss of both H3K23ac and H3K14ac (**Fig. 1C**).

**Figure 1.**
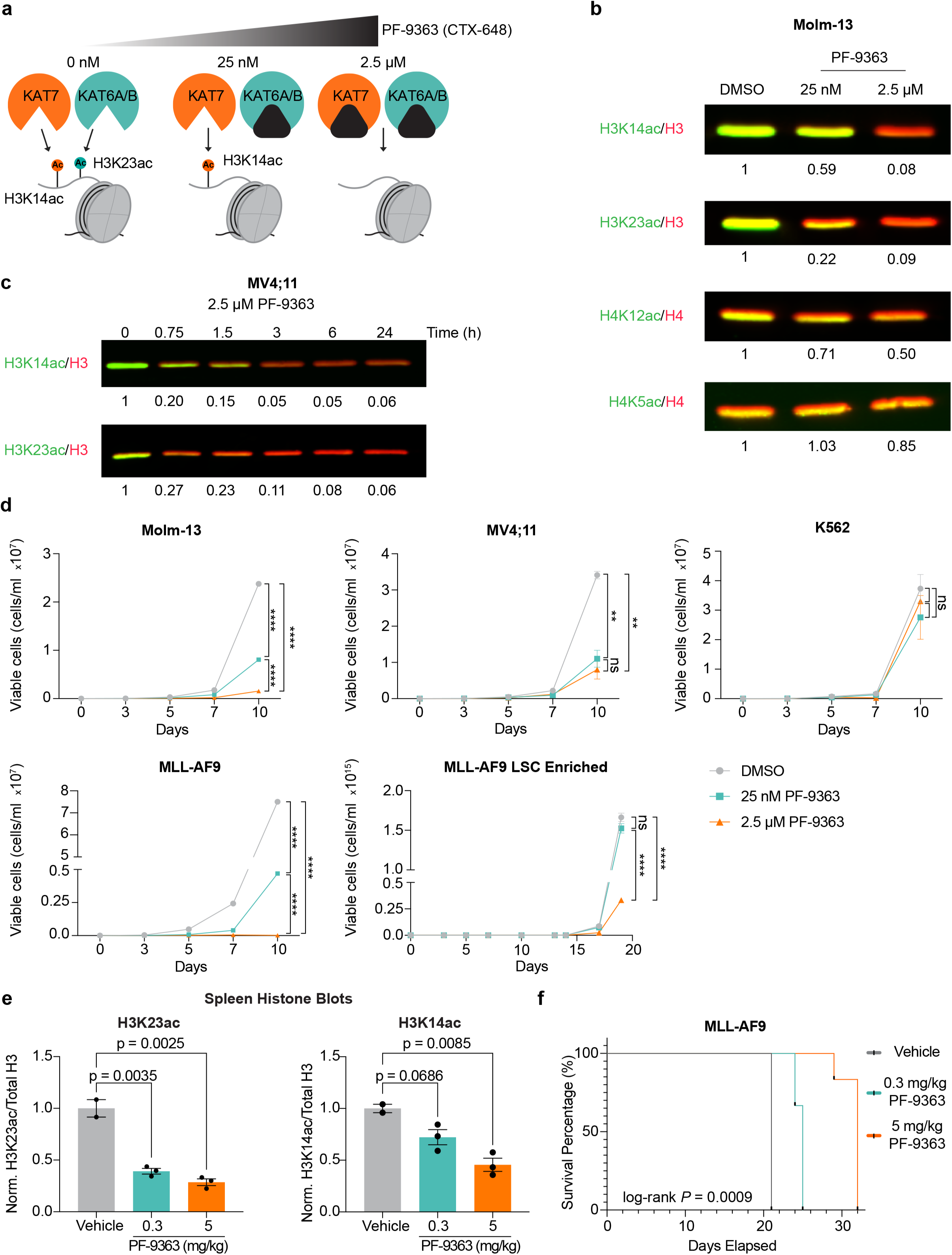
PF-9363 selectively inhibits KAT6A/B and KAT7 to disrupt growth of MLL-fusion driven AML **a,** Schematic depicting doses at which PF-9363 inhibits KAT6A and KAT6B (cyan) and KAT7 (orange). **b,** Histone immunoblots of H3K14ac (KAT7), H3K23ac (KAT6A/B), H4K12ac (KAT7), H4K5ac (KAT5) and total Histone 3 (H3) or total Histone 4 (H4) of Molm-13 cells treated for 24 hr with DMSO, 25 nM or 2.5 µM of PF-9363. Total H3 or H4 blots are shown in red and the relevant histone acetylation mark in green. Values indicate the ratio of the indicated histone mark relative to total H3/H4 abundance, normalised to DMSO. **c**, Time course H3K14ac and H3K23ac immunoblots of MV4;11 cells treated with 2.5 µM of PF- 9363. Total H3 is shown in red and H3K14ac or H3K23ac in green. Values indicate the ratio of H3K14c / H3K23ac mark relative to total H3 abundance, normalised to the 0 hr timepoint. **d**, Proliferation assays of human (Molm-13, MV4;11) and murine (MLL-AF9, MLL-AF9 LSC enriched) leukaemia cell lines and non-AML human cell line K562. Cells were treated with DMSO (grey), 25 nM PF-9363 (cyan) or 2.5 µM (orange) PF-9363. Each dot represents the mean of 3 independent replicates with error bars showing mean ± S.E.M. Statistics were calculated using a two-tailed t-test comparing DMSO versus 25 nM PF-9363 or 2.5 µM PF-9363 and 25 nM PF-9363 versus 2.5 µM PF-9363 at the latest time point indicated on the plot (****p < 0.0001, ***p < 0.001, **p < 0.01, *p < 0.05, ns = p > 0.05). **e**, Bar plot of quantified immunoblots for H3K23ac (left) and H3K14ac (right) from spleens of mice transplanted with MLL-AF9 and treated with vehicle (grey), 0.3 mg/kg PF-9363 (cyan) or 5 mg/kg PF-9363 (orange). Bar plot (mean ± S.E.M) indicates the ratio of H3K14c or H3K23ac mark relative to total H3 abundance, normalised to the vehicle. Statistics were calculated using a two-tailed t-test comparing the means of vehicle versus 0.3 mg/kg PF-9363 or 5 mg/kg PF-9363. **f**, Kaplan-Meier survival curve of NSG mice (n=6 per group) transplanted with MLL-AF9 cells and treated with vehicle (grey), 0.3 mg/kg PF-9363 (cyan) or 5 mg/kg PF-9363 (orange). Cells were injected via tail vein before randomisation into treatment groups and starting dosing 10 days following transplantation. Statistics were calculated using the log-rank (Mantel-Cox) test comparing survival of mice treated with vehicle or 5 mg/kg PF-9363.

Having established the ability to distinguish catalytic inhibition of KAT6A/B from concurrent inhibition of KAT7 we explored the consequences of inhibiting these MYST acetyltransferases in a range of human and murine AML cells driven by different MLL-fusions. These data showed that concentrations of PF-9363 which concurrently inhibited KAT7 more effectively inhibited proliferation than inhibition of KAT6A/B alone (**Fig 1D**). This is particularly pronounced in isogenic MLL-AF9 leukaemia cells which we have previously shown to be enriched for leukaemia stem cell properties(20,52,53). We next wanted to explore the differential effects of inhibiting KAT6A/B alone or in combination with KAT7 in pre-clinical mouse models of MLL-FP leukaemia. To address this question, we initially titrated the effects of PF-9363 drug concentrations (up to the maximum tolerated dose) to establish doses of the drug that primarily inhibited KAT6A/B (0.3 mg/kg) and a higher dose (5 mg/kg) that had more substantial effect on KAT7 (**Fig. 1E**). Using these doses, we treated mice transplanted with the same MLL-AF9 leukaemia cells. These data, which were independently replicated, revealed a clear dose dependent increase in survival emphasising that consistent with our findings *in vitro,* doses of PF-9363 that also inhibit the catalytic activity of KAT7 provide the greatest therapeutic response (**Fig. 1F** and **Fig. S1A**). Taken together these data provide the first *in vivo* evidence that catalytic inhibition of KAT6A/B and KAT7 have a major therapeutic effect in aggressive mouse models of MLL-FP leukaemia. Importantly, these data also emphasise that greater therapeutic efficacy is achieved when KAT7 is inhibited in this disease.

### Systematic analysis of KAT6A, KAT6B and KAT7 in MLL-leukaemia

Whilst we were able to separate the catalytic inhibition of KAT6A/B from KAT7, at present the highly conserved nature of KAT6A and KAT6B cannot be divorced pharmacologically. Moreover, with current chemistry KAT7 cannot be selectively inhibited without concurrent inhibition of KAT6A/B. However, understanding the contribution of each of these three evolutionarily conserved acetyltransferases is important as future advances in medicinal chemistry may offer the opportunity to selectively inhibit these enzymes individually offering the opportunity to maintain the therapeutic effects whilst minimising the side-effects from broader target engagement. Therefore, to systematically address the importance of each of these acetyltransferases in MLL-FP leukaemia, we took a genetic approach using CRISPR/Cas9 mediated knockout of each target individually. Our data showed that whilst genetic loss of KAT6A and to a lesser extent KAT6B had modest effects on leukaemia cell growth, the impact of KAT7 loss was more profound (**Fig. 2A** and **Fig. S1B**). These findings are in line with our previous work showing that conditional knockout of KAT6A in a mouse model of MLL-AF9 leads to prolonged survival; however, genetic loss of KAT7 led to a complete impairment of leukaemia maintenance *in vivo*(20).

**Figure 2.**
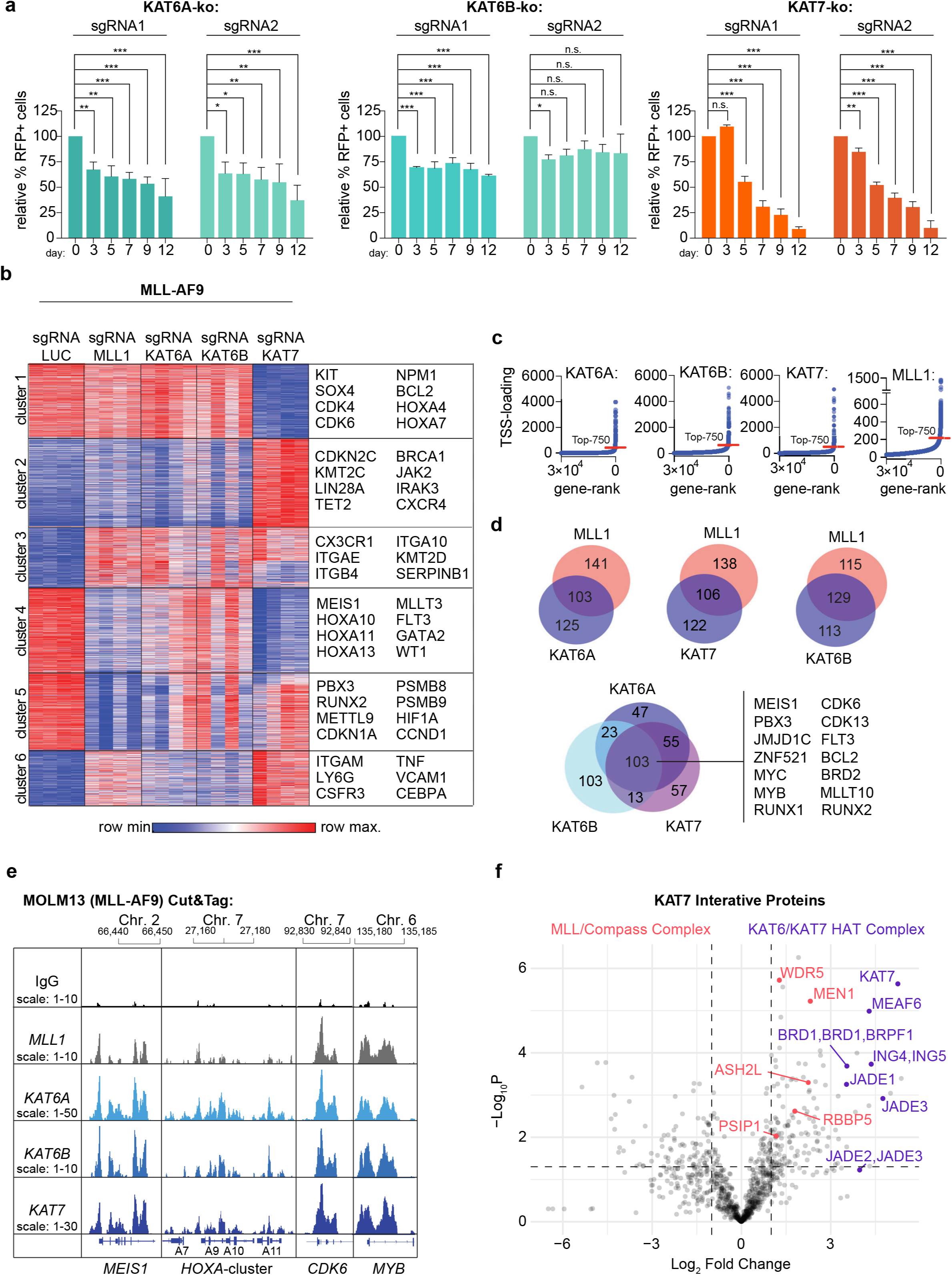
Systematic genetic profiling of MYST KATs reveal KAT7 but not KAT6A/B modulates key targets of MLL-leukaemias **a,** Negative selection competition assays in murine MLL-AF9 cells expressing *S. pyogenes* Cas9 and guide RNAs targeting KAT6A, KAT6B and KAT7. Values represent fraction of RFP-positive CRISPR/Cas9 knockout cells relative to baseline, n=3. Statistics were calculated using an Ordinary One-way ANOVA (*p < 0.05, **p < 0.01, ***p < 0.001). **b**, Heatmap of differentially expressed genes after genetic knockout of Mll1, Kat6a, Kat6b and Kat7 in MLL-AF9 cells. Differentially expressed genes (DESeq2, padj < 0.05, fold-change > 1.5) were clustered by K-means (6) with representative genes from each cluster annotated. **c**, Hockey-stick plots showing asymmetric loading of KAT6A, KAT6B, KAT7 and MLL1 at transcription start sites across the genome in Molm-13 cells. The top 750 TSS were considered as highly loaded sites for the annotation of high-confidence gene targets. **d**, Venn diagram showing overlap of high-confidence gene targets of MLL1 and KAT6A, MLL1 and KAT7 and MLL1 and KAT6B (top). High-confidence KAT6A, KAT6B and KAT7 gene targets are shared at 103 genetic loci including known MLL-AF9 targets (bottom). **e**, Cut&Tag for KAT6A, KAT6B, KAT7 and MLL1 in Molm-13 cells. Representative genome- browser snapshots (IGV) of KAT6A, KAT6B, KAT7 and MLL1 genomic co-localisation at MEIS1, HOXA, CDK6 and MYB. **f**, Volcano plot of enriched proteins in KAT7 immunoprecipitation mass spectrometry versus IgG control in MCF7 cells. Proteins apart of the KAT6A or KAT7 histone acetylation complex are highlighted in purple and proteins apart of the MLL/Compass complex are in pink. Enriched proteins are defined as proteins with and adjusted p-value < 0.05 and an absolute log2 fold change of 1.

To further explore the molecular reasons that underpin these results in MLL-FP leukaemia, we performed RNA-Seq analysis specifically aiming to address the commonalities and differences in the transcriptional programs regulated by KAT6A, KAT6B and KAT7 and relate this to the gene expression regulated by MLL1. These data can be broadly categorised into six clusters (**Fig. 2B**). These include genes that are uniquely regulated by KAT7 (clusters 1 and 2), and genes mainly regulated by MLL1 (cluster 5). The most striking finding from these data is that most of the genes regulated by MLL1 are also affected to some extent by loss of either KAT6A, KAT6B or KAT7 (**Fig. 2B** and **Fig. S1C-E**). Notably, the loss of KAT7 results in more pronounced gene expression changes compared to KAT6A/B and extends the spectrum of affected MLL1-target genes. This is especially prominent with regards to the induction of differentiation (cluster 6) and repression of the homeobox genes, such as MEIS1 and the HOXA cluster (cluster 4), which are central to the pathogenesis of MLL-FP leukaemia (**Fig. 2B** and **Fig. S1C-E**). As several genes were co-regulated by these proteins we next sought to understand if these effects were mediated by co-occupancy of these epigenetic regulators at chromatin. In this regard, our data clearly illustrate that up to half of the transcription start sites (TSS) bound by MLL1 are also bound by KAT6A, KAT6B or KAT7 and the majority of genes are co-occupied by all three MYST acetyltransferases (**Fig. 2C-E**). The co-regulation of a common subset of genes and co-occupancy at chromatin raised the prospect of a dynamic functional interplay between members of these epigenetic complexes. To further understand this functional interaction, we created an isogenic cell line with and without KAT7 (**Fig. S1F**) and used these cells to characterise the KAT7 protein interactome (**Fig. 2F**). These data clearly identified all the members of the KAT7 complex and several interacting partners of the MLL1/Compass-like complex. Notably one of the top protein-protein interactions identified was with Menin.

### KAT7 is a preserved dependency in Menin inhibitor resistant MLL leukaemia cells

Menin is a protein which physically interacts with MLL1(54) and facilitates the chromatin binding to critical MLL1 and MLL-FP target genes(55–60). Importantly, pre-clinical work clearly demonstrated that inhibition of the Menin/MLL interaction led to the loss of expression of key MLL-FP target genes, particularly the homeobox genes, such as MEIS1 and this resulted in impressive efficacy in models of MLL-FP leukaemia(16,17). These seminal observations laid the platform for the clinical trials which have established the importance of Menin inhibition in future clinical management of MLL-FP leukaemia. Despite this substantial progress, the clinical trial data emphasises that although Menin inhibitor monotherapy is effective, genetic resistance due to Menin mutations or non-genetic adaptive mechanisms emerge and re-establish leukaemia(19,61). These findings emphasise the significance of combination therapies and highlight the importance of targeting the gene expression program driven by MLL-FP through diverse and non-redundant targets.

Given the established functional interaction between KAT7 and Menin, we wanted to explore whether targeting KAT6 and/or KAT7 was capable of re-instating therapeutic efficacy in Menin inhibitor resistant leukaemia cells. To address this question, we first established an isogenic model of drug naïve and Menin-inhibitor resistant cells using the OCI-AML2 cell line, which contains the MLL-AF6 oncoprotein. In this model system, resistance was generated by chronic exposure to the Menin inhibitor VTP50469, leading to adaptation and ultimately outgrowth of drug-resistant leukaemia cells (**Fig. S1G**). We sequenced the resistant cells to understand if the resistance phenotype was due to mutations in Menin that are known to hinder drug binding(19). Our data did not reveal any mutations in Menin or other newly acquired somatic mutations that were not already detected in the drug naïve cells (**Fig. S2A**). Of note, a small p53 mutant subclone was selected for in the Menin-inhibitor resistant cells, suggesting a role of p53 in adaptation to Menin inhibitor induced cell death. We then used these cells to assess the effects of CRISPR/Cas9 mediated knockout of either KAT6A, KAT7 or non-catalytic members of these histone acetyltransferase complexes. Surprisingly, we found that cells which were resistant to Menin inhibition also showed resistance to the genetic loss of KAT6A or BRPF1, a member of the KAT6A/B complex (**Fig. 3A**). In contrast, genetic loss of KAT7 and ING5, a member of the KAT7 complex, was equally effective in the drug naïve and isogenic Menin inhibitor resistant cells (**Fig. 3B**).

**Figure 3.**
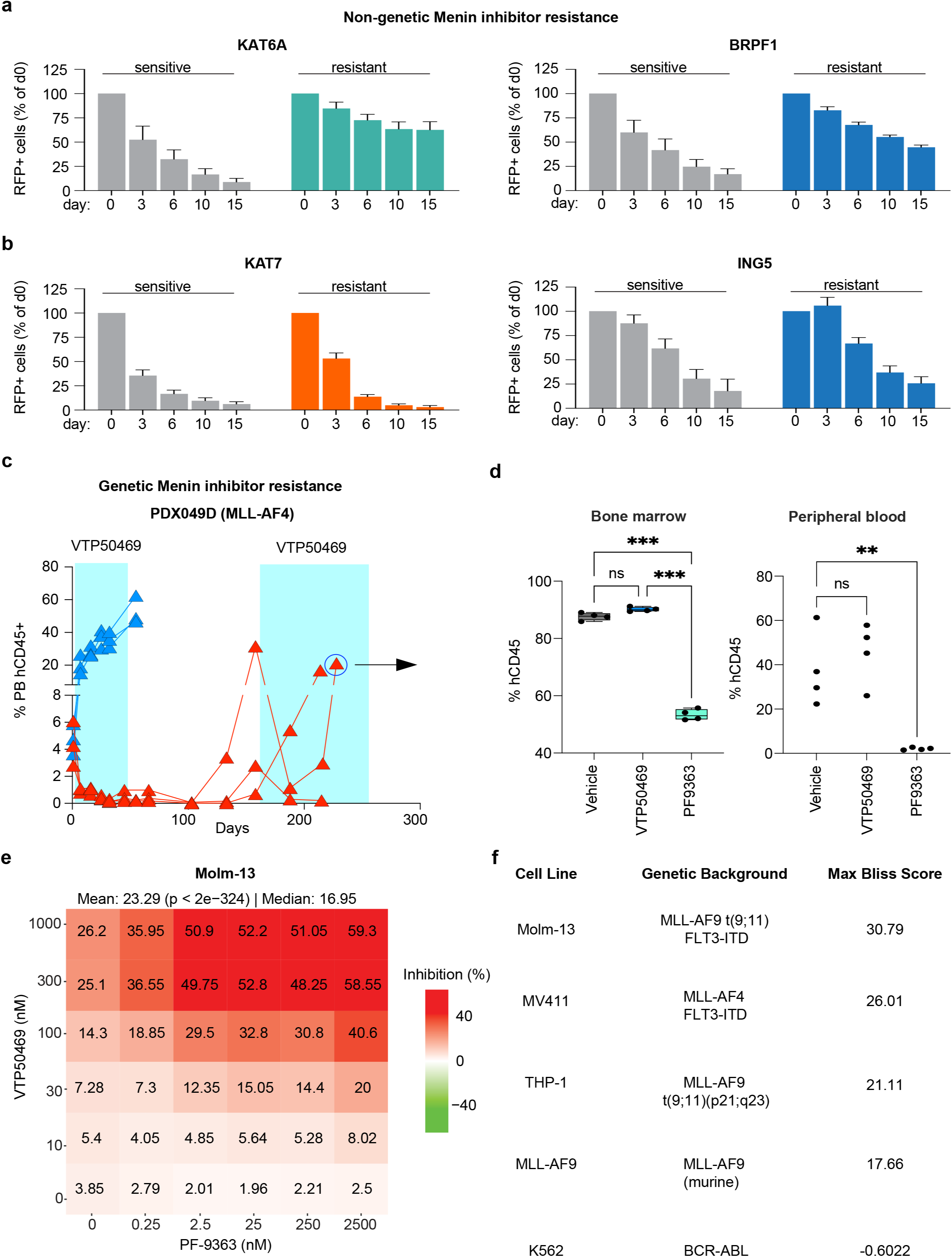
KAT7 is a preserved dependency in Menin inhibitor resistant MLL leukaemia cells. Negative selection competition assays in OCI-AML2 cells resistant and sensitive to Menin inhibition expressing Cas9 and guide RNAs targeting KAT6A or BRPF1 (**a**) and KAT7 or ING5 (**b**). Values represent fraction of RFP-positive CRISPR/Cas9 knockout cells relative to baseline. **c**, Percentage of human CD45-positive cells in peripheral blood of mice transplanted with PDX049D and treated with 0.03% VTP50469 (in rodent diet). **d**, Percentage of human CD45-positive cells in the bone marrow and peripheral blood of NOG mice transplanted with Menin resistant PDX049D treated with vehicle, 0.1% VTP50469 (in rodent diet) or 5 mg/kg PF-9363 (intraperitoneal injection) for 14 days. **e**, Molm-13 cells were treated for 96 hr with indicated doses of PF-9363 and/or VTP50469. Values indicate the percentage of non-viable cells relative to indicated doses of drug. The mean and median of the percent non-viable cells across all conditions are displayed above the heatmap. **f**, Summary table of Bliss synergy scores between PF-9363 and VTP50469 for indicated leukaemia cell lines, Molm-13, MV4;11, THP-1 and MLL-AF9. Synergy scores were calculated using the Bliss method. Max synergy scores are defined as the Bliss synergy score that corresponds to the combination of the highest concentration of both drugs, 1 µM VTP50469 and 2.5 µM PF-9363.

We next wanted to address the possibility that targeting KAT6/KAT7 may also overcome genetic resistance to Menin inhibitors driven by target site mutations in the *MEN1* gene. Here we used a patient derived xenograft (PDX) model which was first treated with the Menin inhibitor in isolation. After cessation of therapy, the cancer re-emerged in several mice and these mice were re-challenged with Menin inhibitor to isolate a population of malignant cells that failed to respond to re-challenge (**Fig. 3C**). We then sequenced these tumours and showed that the resistance in these mice were due to the presence of mutations in Menin (G133D and T349M) which we had previously shown to negate drug binding(19) (**Fig. S2B**). The *MEN1* mutated leukaemia was then re-transplanted into secondary recipient mice and following engraftment challenged with either the Menin inhibitor (VTP50469) or PF-9363. These data clearly show that catalytic inhibition of KAT6/KAT7 is able to overcome genetic resistance to the Menin inhibitor (**Fig. 3D**).

Whilst these data provide strong genetic and pharmacological evidence to support the notion that targeting of KAT6/KAT7 could overcome genetic and non-genetic mechanisms of resistance to Menin inhibitors, it was unclear if these epigenetic regulators functioned in an epistatic manner or if they have a non-redundant relationship which could be exploited with combination therapies to enhance the efficacy of these drugs. To answer this important question, we performed synergy assays using PF-9363 and VTP50469 in a range of human and murine leukaemia cells containing an MLL-FP. Our findings demonstrate marked synergy between PF-9363 and VTP50469 in all AML cells containing an MLL-FP (**Fig. 3E-F** and **Fig. S3A-H**), but these effects were not observed in leukaemia cells driven by an alternative fusion oncoprotein such as BCR-ABL (**Fig. 3F, S3D, H**). Notably, as exemplified by the data in Molm-13 cells (**Fig. 3E**), the efficacy of VTP50469 at a moderately effective dose (100 nM) is enhanced by selective inhibition of KAT6A/B with PF- 9363 (25 nM) but this effect is more markedly potentiated when KAT7 is inhibited with PF-9363 (2.5 µM). PF-8144 is a recently described clinically active inhibitor(62) with a similar selectivity profile to PF-9363. Treatment of MLL-FP leukaemia cells with this drug, similarly, shows a spectrum of sensitivity with some cell lines showing a response at concentrations that primarily target KAT6A/B whilst several require higher concentrations that concomitantly inhibit KAT7 to achieve maximum efficacy (**Fig. S4A**). Notably, the responsiveness of the cells to drug is not correlated with the expression levels of KAT6A, KAT6B or KAT7 (**Fig. S4B-E**). Importantly, even with this chemically distinct compound we see marked synergistic efficacy when combined with a Menin inhibitor resulting in more dramatic differentiation and cell death (**Fig. S4F-I**).

### Dual inhibition of Menin and KAT7 results in a rapid and marked repression of MLL-FP target genes

Our genetic and pharmacological experiments provided orthogonal evidence to support the importance of concurrently targeting KAT6/KAT7 with Menin inhibition in MLL-FP. Therefore, we next sought to understand the molecular basis for this strong synergistic benefit. To initially study the gene expression changes induced by PF-9363 and VTP50469 in isolation and in combination we performed RNA-seq analyses in Molm-13 cells, a human leukaemia cell line driven by MLL-AF9. Here we initially assessed the gene expression changes with these inhibitors at an early time point (t = 6 hours). We chose 6 hours as this is a time frame that allows for the decline in steady state mRNA and enables a clear identification of early changes in gene expression. This time frame is particularly important as the reported median half-life of the clinical Menin inhibitor is ∼3 hours(18), so we also wanted to assess how a time-limited clinical exposure to effective on-target inhibition alters gene-expression in the MLL-FP leukaemia cells.

Our data show that at this early time point there is only very modest changes in gene expression with single agent therapies (**Fig. 4A-B** and **Fig. S5A**); however, combination therapy results in a marked repression of the key genes central to the MLL-FP leukaemia transcription program including the homeobox genes MEIS1, HOXA9 and PBX3 (**Fig. 4C-D**). Notably, the gene expression changes with PF-9363 have some shared but several distinct features when compared to VTP50469 treatment. Most striking is that fact that combination therapy clearly enhanced repression of both the shared and distinct gene sets from the individual treatments resulting in a composite signature associated with myeloid differentiation (**Fig. 4E**) and a prominent repression of genes previously identified to be directly bound by the MLL-AF9 fusion in mouse leukaemia cells(55) (**Fig. 4F**). The gene expression changes seen with the combination therapy within 6 hours is close to the maximal effect, as a longer duration of combined inhibition for up to 24 hours does not result in an appreciable change in gene repression (**Fig. S5B-E**). To further illustrate the molecular consequences of the combination therapy we performed ChIP-seq which demonstrates a near complete loss in chromatin occupancy for MLL1 and Menin at the genes repressed by the combination therapy (**Fig. 4G**).

**Figure 4.**
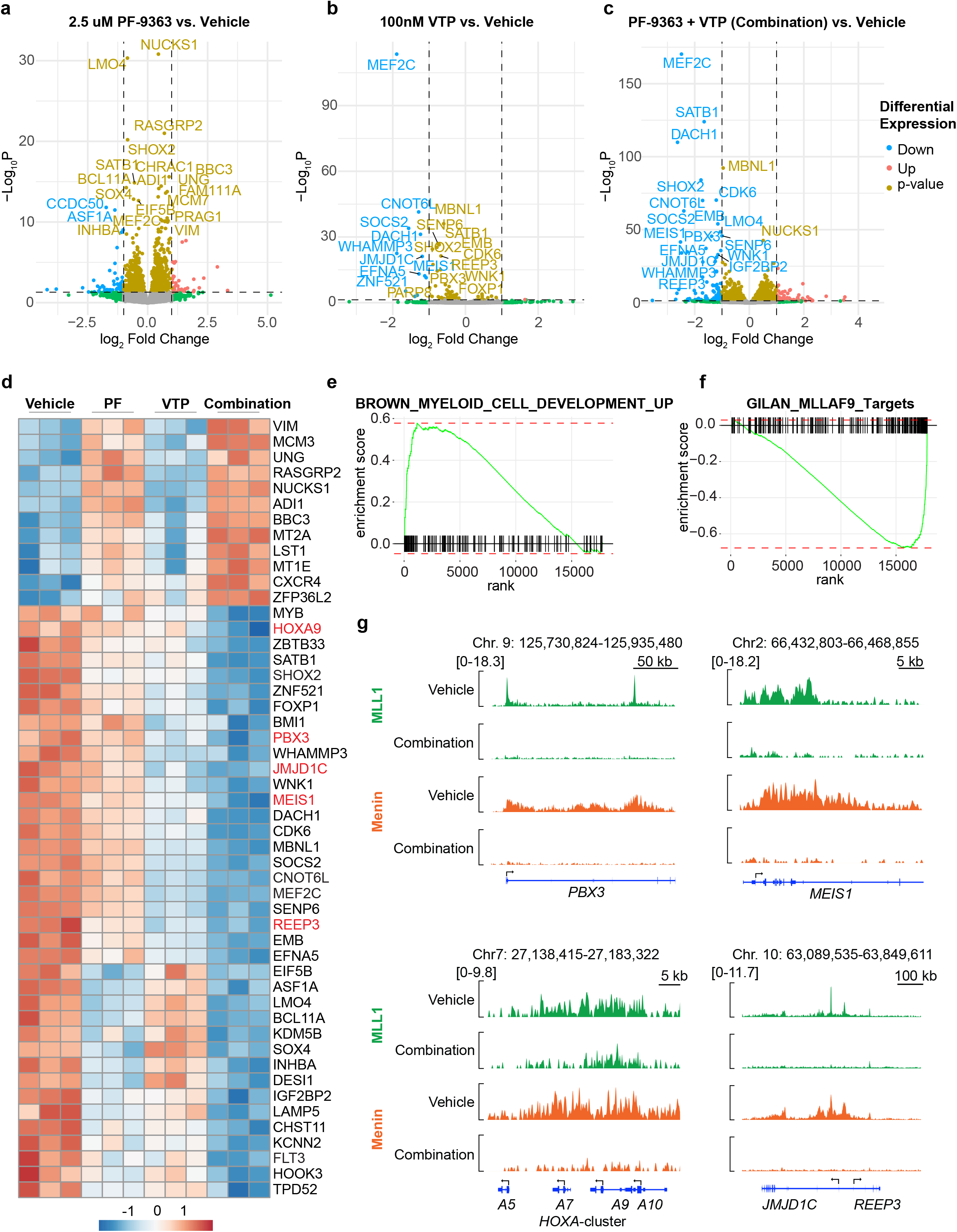
Combination of PF-9363 and VTP50469 induces rapid shutdown of the leukemic transcriptional program Volcano plot showing differential gene expression of Molm-13 cells treated for 6 hr with 2.5 µM PF-9363 versus vehicle (**a**), 100 nM VTP50469 versus vehicle (**b**) or combination (2.5 µM PF- 9363 and 100 nM VTP50469) versus vehicle (**c**). Significant differentially expressed genes are defined as genes with an adjusted p-value < 0.05 and absolute log2 fold change of 1. **d**, Heatmap showing the top 50 differentially expressed genes by adjusted p-value for the combination versus vehicle treatment groups after 6 hr of treatment. Key targets of the MLL-AF9 oncogene, MEIS1 PBX3, HOXA9, REEP3 and JMJD1C are highlighted in red. Row scaled Z-scores are shown. PF = 2.5 µM PF-9363 treatment group, VTP = 100 nM VTP50469 treatment group. GSEA of the BROWN_MYELOID_CELL_DEVELOPMENT gene set **(e)** and the GILAN_MLLAF9_Targets gene set (**f**) for the combination versus vehicle treatment groups. Positive scores indicate enrichment in the combination group. Negative scores indicate enrichment in the vehicle group. **g**, Chromatin immunoprecipitation for MLL1 and Menin in Molm-13 cells treated for 24 hr with vehicle or combination (2.5 µM PF-9363 and 100 nM VTP50469). Genome browser snapshots of *PBX3*, *HOXA* cluster, *MEIS1, REEP3* and *JMJD1C*.

### Synergistic benefit of PF-9363 and VTP50469 in MLL-FP leukaemia

Having established the molecular rationale for combination therapy with PF-9363 and VTP50469, we next wanted to explore whether the more profound transcriptional inhibition of MLL-FP leukaemia genes seen with this combination also resulted in a greater pre-clinical efficacy. To approximate the clinical scenario of primary resistance to Menin inhibition, we chose a PDX model of MLL-AF6 AML which showed only a modest response to single agent Revumenib (SNDX- 5613) and single agent PF-9363 after 28 days of treatment (**Fig. 5A** and **Fig. S5F-H**). In contrast, treatment with combination therapy induced a more marked reduction in leukaemia burden in the bone marrow, peripheral blood and spleen (**Fig. 5A**). The combination therapy was not associated with greater cytopenia (**Fig. S5H**) but induced greater differentiation of the leukaemia cells (**Fig. S5F**). Whilst PDX models provide a closer approximation to the clinical scenario, technical limitations often preclude the molecular insights that can be gleaned to explain these promising pre-clinical results. Therefore, to better study the molecular consequences of these therapies at clonal resolution *in vivo* we employed **SPLINTR** (**S**ingle-cell **P**rofiling and **LIN**eage **Tr**acing)(63) in a xenograft model of Molm-13 leukaemia (**Fig. S6A**). SPLINTR employs unique heritable DNA barcodes that are transcribed and can be readout using various single-cell platforms to link clonal fate to gene expression in a temporally resolved manner at single clone resolution. Moreover, to distinguish cell-intrinsic and cell-extrinsic effects on response to these therapies we used a clone-splitting approach(63). In this strategy, every mouse receives an identical population of malignant clones; consequently, if the same clone exhibits the same behaviour (e.g. resistance) in all mice, then it is clear that a cell-intrinsic state underpins its adaptive potential.

**Figure 5.**
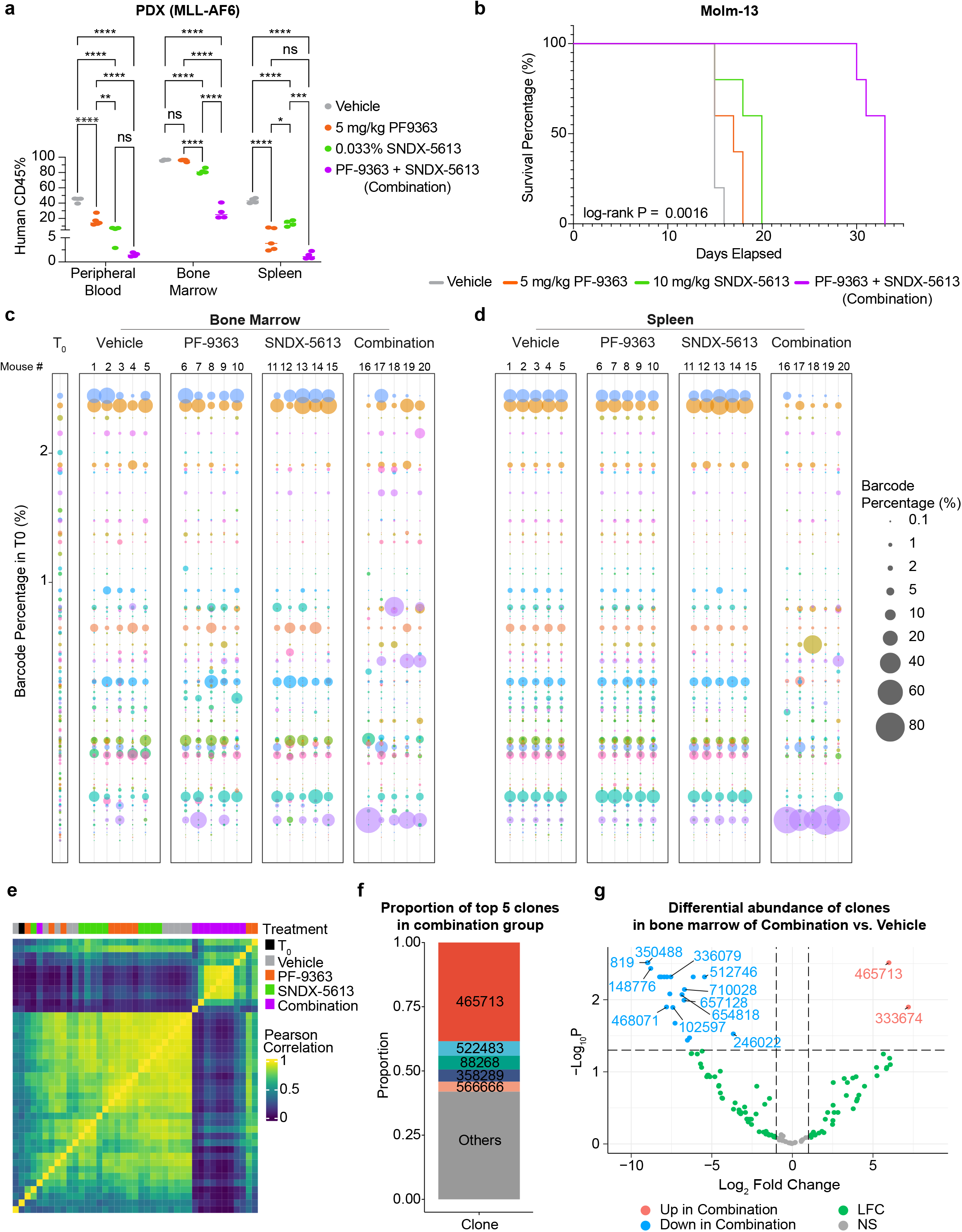
Combination therapy extends disease latency by restricting outgrowth of leukemic clones **a**, Percentage of human CD45-positive cells in peripheral blood, bone marrow and spleen of mice transplanted with PDX (MLL-AF6) and treated for 28 days with vehicle (grey), 5 mg/kg PF-9363 (orange), 0.033% SNDX-5613 (green) or combination (5 mg/kg PF-9363 and 0.033% SNDX- 5613, purple). Statistics were calculated using a two-tailed t-test comparing means of each treatment group against each other (****p < 0.0001, ***p < 0.001, **p < 0.01, *p < 0.05, ns = p > 0.05). **b**, Kaplan-Meier survival curve of NSG mice transplanted with Molm-13 cells and treated with vehicle (grey), 5 mg/kg PF-9363 (orange), 10 mg/kg SNDX-5613 (green) or combination (5 mg/kg PF-9363 and 10 mg/kg SNDX-5613, purple). Dosing commenced 7 days after transplantation and continued for the duration of the experiment. Log-rank (Mantel-Cox) test comparing survival of mice in the vehicle and combination treatment groups. **c**,**d** SPLINTR barcode-sequencing data from baseline (T0) and experimental endpoint for each treatment group (Vehicle, PF-9363, SNDX-5613 and Combination) in the bone marrow (**c**) and spleen (**d**). Clones are ranked by their percentage in the baseline with each coloured circle representing a unique clone. **e**, Heat map of pairwise Pearson correlation values for barcode-sequencing samples from each treatment group T0 (black), vehicle (grey), PF-9363 (orange), SNDX-5613 (green) and combination (purple). **f**, Stacked bar plot showing the proportion of the 5 largest barcoded clones in the combination group compared to the proportion of all other clones. Others group indicates proportion of all other clones. **g**, Volcano plot showing differential barcode abundance of clones in bone marrow of mice treated with combination versus vehicle. Adjusted p-value < 0.05 and absolute log2 fold change of 1.

We specifically chose the Molm-13 xenograft model as we found that doses of PF-9363 (**Fig. 1E**) and VTP50469(17), which showed a clear survival advantage in other models of MLL-AF9 leukaemia but failed to result in a survival benefit in this highly aggressive leukaemia model consequently approximating the PDX model and clinical scenario of primary resistance (**Fig. 5B** and **Fig. S6B**). Importantly and in contrast to the single agent therapies, we consistently found that combination therapy resulted in a marked survival benefit (**Fig. 5B** and **Fig. S6B**). Our clonal analysis revealed that leukaemia initiating potential was highly cell-intrinsic with exactly the same clones giving rise to leukaemia in both medullary (bone marrow) and extra-medullary (spleen) sites (**Fig. 5C-D**). Interestingly, ongoing treatment with either single agent PF-9363 or VTP50469 did not alter the clonal structure or clonal composition of the leukaemia at the time of fulminant disease, showing high correlation in all mice treated with either vehicle or single agent therapy (**Fig. 5C-E**).

In contrast, combination therapy substantially altered clonal structure whereby only a minority of clones were able to withstand the increased pressure exerted by the combination (**Fig. 5C-E** and **Fig. S6C**). Notably, the clones that survived the combination therapy were not the major leukaemia initiating clones that established the disease in the absence of treatment or indeed the clones most adept at surviving monotherapy with either drug (**Fig. 5E**). In fact, approximately 60% of the overall resistant leukaemia burden in all the mice treated with combination therapy was composed of just 5 clones (**Fig. 5F**). Notably, a single clone (465713), was the dominant non-responsive clone comprising >40% of the overall disease burden in both the bone marrow and spleen samples (**Fig. 5F**).

### Resistant clones do not reinstate the canonical MLL-FP transcription program and instead adapt to express SLPI

Although just 5 clones contributed to majority of the resistant leukaemia in all mice treated with combination therapy, several smaller clones were also noted to be preferentially enriched in the leukaemia cells after combination therapy (**Fig. 5F-G** and **Fig. S6D-E**). To understand the differential molecular features that distinguished the responsive and non-responsive clones we coupled our SPLINTR barcoding approach with single cell RNA sequencing. Here we noted that the canonical MLL-FP target genes such as HOXA9 and MEIS1 were highly expressed in the leukaemia cells prior to transplantation and that this expression was preserved in the cells that engrafted and gave rise to fulminant leukaemia in the vehicle treated mice *in vivo* (**Fig. 6A-B** and **Fig. S6F**). Surprisingly, the leukaemia cells that were resistant to combination therapy with PF- 9363 and VTP5046 do not re-instate the canonical MLL-FP transcriptional at the time of death from resistant disease (**Fig. 6A-B**). These resistant cells continue to show a strong repression of HOXA9 and MEIS1 and instead show a marked enrichment for various inflammatory gene expression programs (**Fig. 6C** and **Fig. S6G-I**).

**Figure 6.**
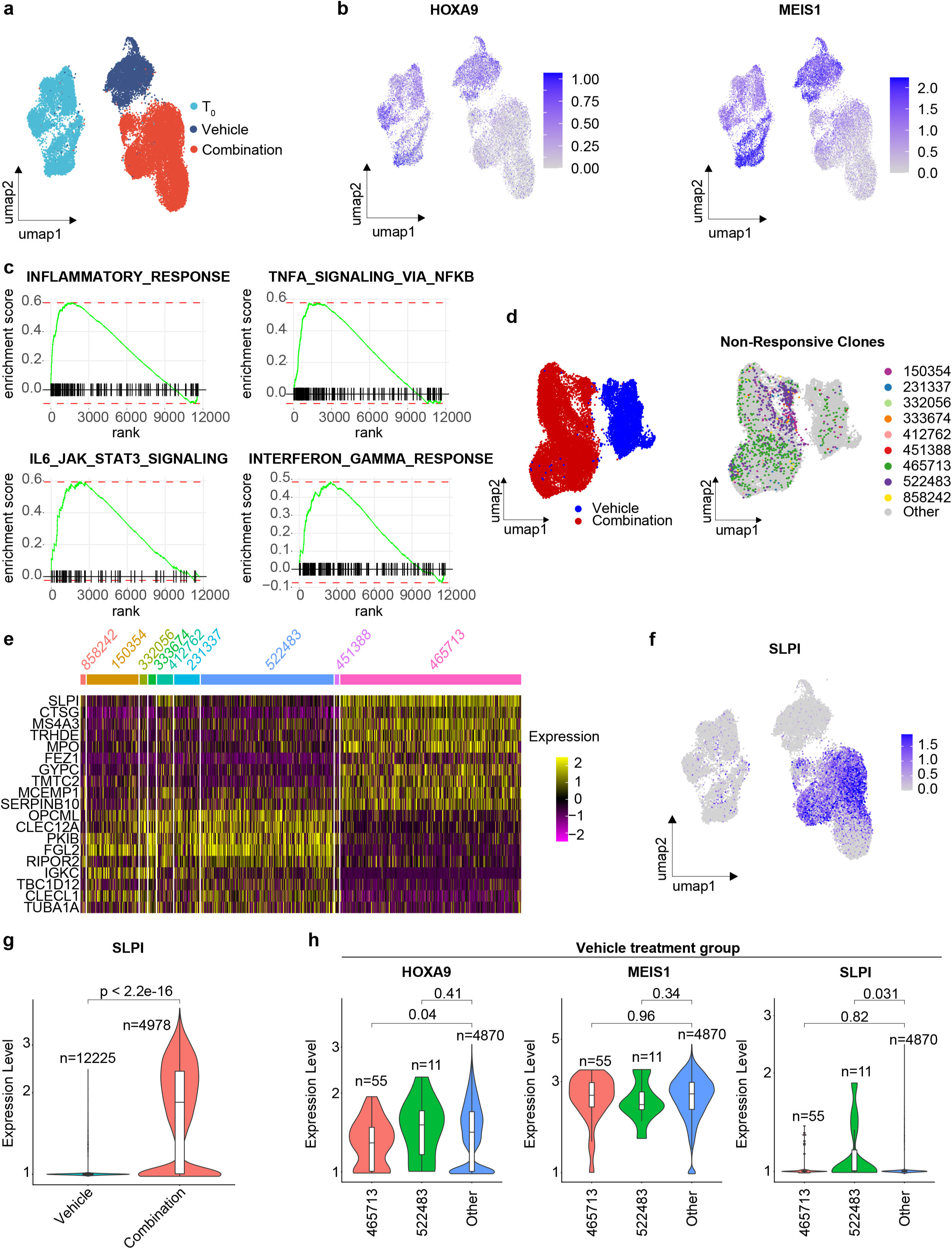
Single cell RNA-Seq reveals clone dependent responses to combined PF-9363 and VTP50469 **a,** UMAP of scRNA-sequencing dataset composed of samples from T0 (light blue), vehicle (dark blue) and combination (red) groups from 24,906 cells. **b**, UMAP annotated with the normalised expression of HOXA9 and MEIS1 in T0, vehicle and combination groups. **c**, GSEA plots of Inflammatory Response, TNF-alpha signalling through NFKB, IL6_JAK_STAT3 signalling, and Interferon Gamma Responsive gene sets in the combination group versus the vehicle group. Positive scores indicate enrichment in the combination group. Negative scores indicate enrichment in the vehicle group. **d**, UMAP of vehicle and combination treatment groups (left) and UMAP annotated for cells comprising non-responsive clones (right). **e**, Heatmap of differentially expressed genes identified from pairwise comparisons of non-responsive clones in the combination treatment group. Heatmap shows row-scaled normalised expression values. **f**, UMAP annotated with the normalised expression of SLPI across T0, vehicle and combination groups. **g**, Violin plot of SLPI expression in vehicle versus combination treatment groups. Wilcox test. **h**, Violin plots of HOXA9 (left), MEIS1 (centre) and SLPI (right) expression in the vehicle treatment group for non-responsive clones 465713 and 522483. Other clones represent clones responsive to combination treatment. Wilcox test.

As SPLINTR enables us to assess the single cell transcriptome at clonal resolution we focused on understanding the specific transcriptional changes seen in the non-responsive clones which are able to withstand the combined pressure exerted by KAT6/7 and Menin inhibition. Our scRNA-seq confirmed our results from the DNA barcoding (**Fig. 5E-G**) showing that resistant clones are overall underrepresented in the vehicle treated leukaemic population (**Fig. 6D** and **Fig. S6G**). When the transcriptome of these clones were analysed, we noted that although these resistant clones do not express a uniform transcription program, there are several genes that are commonly expressed within multiple clones (**Fig. 6E**). Remarkably, the most prominent gene expressed across multiple resistant clones is SLPI (secreted leucocyte peptidase inhibitor), a prominent mediator implicated in altering the tumour microenvironment(64,65), and importantly a gene which we recently identified and functionally validated to be a major driver of malignant clonal fitness in AML including MLL-FP leukaemia(63). Although SLPI is markedly overexpressed in the resistant population as a whole (**Fig. 6F-G**), we have the ability to assess gene expression in identical clones prior to and following treatment to resolve whether this prominent expression of SLPI was pre-existing or emerged as an adaptive process to therapeutic challenge. When we compared the expression of SLPI, MEIS1 and HOXA9 in our two dominant non-responsive clones (465713 and 522483) we found that in the absence of treatment these clones highly express HOXA9 and MEIS1 but not SLPI (**Fig. 6H**). Interestingly, although the level of expression of SLPI in clone 522483 in vehicle treated mice is variable, the dominant clone 465713 which has the highest average expression of SLPI following combination treatment does not express SLPI without therapeutic challenge (**Fig. 6H**). This finding clearly highlights the power of SPLINTR in understanding malignant behaviour and strongly suggests that rather than selecting clones with a pre-existing signature associated with resistance, the resistant clones which survived combination treatment were the clones most adept at rapidly adapting their transcriptional program to negotiate the potent therapeutic challenge.

Together, our findings highlight several critical points; firstly, the primary resistance to Menin inhibitors seen in some MLL-FP leukaemias can be substantially overcome with concurrent dual catalytic inhibition of KAT6/KAT7, which results in a more profound and broader suppression of MLL-FP transcription program. The selective pressure exerted by this combined inhibition can only be overcome by rare malignant clones that are able to rapidly adapt to the therapeutic challenge to survive without the canonical MLL-FP gene expression program. Finally, these data also reinforce the importance of prominent mediators such as SLPI which implicated in altering the tumour microenvironment(63–65) to facilitate malignant fitness.

## DISCUSSION

MLL-FP leukaemia remains an aggressive and often incurable disease. However, decades long investment in studying the molecular pathogenesis of this cancer has yielded several novel therapeutics. Whilst a broad range of candidates aimed at negating the transcriptional programs that sustain this leukaemia have entered early clinical trials(15), the therapy with the most promising clinical results have been small molecule inhibitors that disrupt the interaction between Menin and MLL1 resulting in the displacement of the oncoprotein from its target genes(16–19). Interestingly, the eviction of the MLL-FP from chromatin by Menin inhibition also results in a marked reduction of H3K79me2 at MLL-FP target genes as DOT1L is concurrently dislocated(16,17,55). Consequently, Menin inhibition, at least in part, phenocopies the gene expression effects of DOT1L inhibition albeit with much more rapid kinetics(55,56).

Menin inhibition alone is unable to provide lasting clinical benefit in most patients, highlighting the pressing need to combine these drugs with other effective agents in MLL-FP leukaemia. Here we established that catalytic inhibitors of the MYST acetyltransferases KAT6A/B and KAT7 provide marked synergistic efficacy in combination with Menin inhibitors. Importantly, we establish that although KAT6A/B inhibition alone is effective in some MLL-FP leukaemia cells, to deliver maximal therapeutic benefit, KAT7 also needs to be effectively targeted. The synergistic efficacy with Menin inhibition is underpinned by a rapid eviction of the MLL-FP from chromatin resulting in a major repression of the transcriptional program driven by this oncoprotein. The fact that even steady state mRNA levels are dramatically reduced within 6 hours of exposure to combination therapy is clinically important, as peak drug concentrations with intermittent dosing is only sustained for a few hours and the overt reliance of MLL-FP leukaemia cells on this transcription program provides an important therapeutic window to target these malignant cells. Here we have specifically focused on MLL-FP as we wanted to provide a clear molecular rationale for the combination therapy. However, it is increasingly clear that disrupting the Menin-MLL interaction has clinical impact beyond MLL-FP disease and other leukaemias including those with the NPM1c mutation that express canonical MLL1 target genes such as HOXA9 and MEIS1 also derive substantial benefit from Menin inhibitors(66,67). As KAT7 also physically associates with wildtype MLL1, future work will evaluate the strong possibility that the molecular and therapeutic effects described here extend more broadly in other subsets of AML.

Our genetic and pharmacological data showed that targeting KAT7 provided an orthogonal approach to inhibiting the critical MLL-FP transcription program with Menin inhibitors. Using leukaemia models of genetic and non-genetic acquired (**Fig. 3A-D**) and primary (**Fig. 5A-B and Fig. S6B**) resistance to Menin inhibition, we highlighted the importance of this non-redundant combination approach in these clinical scenarios. Although current chemistry is unable to selectively divorce catalytic inhibition of KAT7 from KAT6A/B, our genetic data strongly advocates for ongoing ingenuity in medicinal chemistry to develop the capacity to selectively inhibit the catalytic activity of KAT7. It is possible that these compounds may maintain the efficacy of the current broader spectrum MYST inhibitors whilst reducing unwanted side-effects. PF-8144, the clinical inhibitor phenocopies our results with PF-9363 (**Fig. S4**). Importantly, this drug this currently under evaluation in breast cancer. These data raise the exciting prospect that our findings of combined targeting of KAT6/KAT7 with Menin in MLL-FP leukaemia could be rapidly translated to further improve the clinical outcomes for what remains a poor prognostic disease.

## ACKNOWLEDGMENTS

We thank members of the Dawson and Armstrong Laboratories for critical review of the manuscript. The Victorian Centre for Functional Genomics (VCFG), The Translational Research Centres, The Molecular Genomics Core and The Research Flow Core at Peter MacCallum Cancer Centre for helpful discussions and technical assistance. We also thank Dr. Jerry McGeehan from Syndax for providing VTP50469 and for valuable discussions. **Funding:** We gratefully acknowledge the following funders for fellowship and grant support: CCV Dunlop Fellowship, NHMRC Investigator grant (1196749), MRFF Research Grant (**1202192**), HHMI_IRS (MAD) and the Peter MacCallum Foundation. SAA is supported by NCI grants CA206963 and CA066996. FP is supported by the Emmy-Noether Programme of the German Research Foundation (DFG, PE 3217/2-1, PN: 528168324), the DFG Research-Unit TARGET-MPN (PE 3217/4-1, PN: 517204983) and the “Else Kröner-Fresenius-Stiftung” (2021-EKEA.111).

## AUTHOR CONTRIBUTIONS

SJVG, FP, SAA & MAD conceptualised the research, designed and analysed experiments and wrote the manuscript with helpful contributions from all authors. JH, EYNL, Y-CC, AG, HH & DV performed computational analyses. SJVG, FP, LM, JP, TK, DVW, WB, JC, JM, BB, MAC, YEB, KK, JB, ENW, DV, JL, SU & JW performed *in vitro* and *in vivo* biology experiments, analysed and interpreted data. PAS, IPS, BJM, SS, TAP were involved in the design and synthesis of the compounds, generated and analysed data. SAA and MAD jointly supervised the work.

## DATA AVAILABILITY

All the sequencing data have been deposited to the NCBI Sequence Archive under the GEO accession number GSE277100.

