## Supplemental Figure Legends for "Catalytic inhibition of KAT6/KAT7 enhances the efficacy and overcomes primary and acquired resistance to Menin inhibitors in MLL leukaemia"

**Supplemental Figure 1**

**a**, Kaplan-Meier survival curve of NSG mice (n=6 per group) transplanted with MLL-AF9 cells and treated with vehicle (grey), 0.3 mg/kg PF-9363 (cyan) or 5 mg/kg PF-9363 (orange). Cells were injected via tail vein before randomisation into treatment groups and starting dosing 10 days following transplantation. Statistics were calculated using the log-rank (Mantel-Cox) test comparing survival of mice treated with vehicle or 5 mg/kg PF-9363.

**b**, Immunoblots of MLL-AF9 cells infected with a control guide or guide RNAs targeting KAT6A or KAT7.

Volcano plots of differentially expressed genes in KAT6A (**c**) KAT6B (**d**) and KAT7 (**e**) knockout MLL-AF9 cells annotated with key targets of the MLL-AF9 oncogene.

**f**, Immunoblots of MCF7 cells infected with a control guide or guide RNAs targeting KAT7. **g**, Dose-response assay showing cell counts (relative to DMSO control) of OCI-AML2 cells resistant and sensitive to menin inhibitor treatment treated with increasing doses of VTP50469.

**Supplemental Figure 2**

**a**, Table of variant allele frequencies detected in drug naïve and drug-resistant OCI-AML2 cell lines from a targeted sequencing panel.

**b**, Table of variant allele frequencies detected in drug naïve and drug-resistant patient derived xenograft, PDX049D, from a targeted sequencing panel.

**Supplemental Figure 3**

MLL-AF9 (**a**), MV4;11 (**b**), THP1 (**c**) and K562 (**d**) cells were treated for 96 hr with indicated doses of PF-9363 and/or VTP50469. Values indicate the percentage of non-viable cells relative to indicated doses of drug. The mean and median of the percent non-viable cells across all conditions are displayed above the heatmap.

MLL-AF9 (**e**), MV4;11 (**f**), THP1 (**g**) and K562 (**h**) cells were treated for 96 hr with indicated doses of PF-9363 and/or VTP50469. Values indicate the percentage of cells that have not undergone growth arrest relative to indicated doses of drug and normalised to the DMSO sample.

**Supplemental Figure 4**

**a,** Half-maximal inhibitory concentration (IC50) and area under the curve (AUC) of PF-8144 across MLL-FP cell lines, MV4;11 (MLL-AF4), THP1, Molm-13, Molm-14, NOMO1 (MLL-AF9), OCI-AML2, ML2 (MLL-AF6), OCI-AML4 (MLL-ENL) and EOL1 (MLL-PTD).

**b**, Expression of KAT6A (left), KAT6B (middle) and KAT7 (right) in MLL-FP cell lines with IC50 values for PF-8144 treatment lower than 100 nM and greater than 100 nM.

Expression of KAT6A (**c**), KAT6B (**d**) and KAT7 (**e**) in MLL-FP cell lines relative to PF-8144 IC50 concentration.

**f**, Experimental schematic of SNDX-5613 and PF-8144 synergy assay.

**g**, OCI-AML2 cells were treated for 7 days with indicated doses of PF-8144 and/or SNDX-5613. Values indicate the Bliss synergy score relative to indicated doses of drug. The mean Bliss synergy score is displayed above the plot.

**h**, Histogram of CD11b fluorescence levels relative to OCI-AML2 cells treated with DMSO (red), 60 nM PF-8144 (blue), 100 nM SNDX-5613 (orange) and combination (60 nM PF-8144 and 100 nM SNDX-5613, green).

**i**, Density plot of CD11b fluorescence levels in OCI-AML2 cells treated with DMSO, 60 nM PF-8144, 100 nM SNDX-5613 and combination (60 nM PF-8144 and 100 nM SNDX-5613).

**Supplemental Figure 5**

**a**, Relative expression of *HOXA9* (right) and *MEIS1* (left) in Molm-13 cells treated for 6, 24, 48 and 72 hr with vehicle (grey), 2.5 μM PF-9363 (orange), 100 nM VTP (green) and combination of 2.5 μM PF-9363 and 100 nM VTP (purple).

Volcano plot showing differential gene expression of Molm-13 cells treated for 24 hr with 2.5 μM PF-9363 versus vehicle (**b**), 100 nM VTP50469 versus vehicle (**c**) or combination (2.5 μM PF-9363 and 100 nM VTP50469) versus vehicle (**d**). Significant differentially expressed genes are defined as genes with an adjusted p-value > 0.05 and an absolute log_2_ fold change of 1.

**e**, Heatmap showing the top 50 differentially expressed genes by adjusted p-value for the combination versus vehicle treatment groups after 24 hr of treatment. Row scaled Z-scores are shown. PF = 2.5 μM PF-9363 treatment group, VTP = 100 nM VTP50469 treatment group.

**f**, Mean fluorescence intensity of CD11b, CD13 and CD14 positive cells in mice transplanted with PDX (MLL-AF6) and treated for 28 days with vehicle (grey), 5 mg/kg PF-9363 (orange), 0.033% SNDX-5613 (green) or combination (combo, 5 mg/kg PF-9363 and 0.033% SNDX-5613, purple).

**g**, Weight (mg) of spleens from mice transplanted with PDX (MLL-AF6) and treated for 28 days with vehicle (grey), 5 mg/kg PF-9363 (orange), 0.033% SNDX-5613 (green) or combination (combo, 5 mg/kg PF-9363 and 0.033% SNDX-5613, purple).

**h**, Complete blood counts (CBC) of mice transplanted with PDX (MLL-AF6) and treated for 28 days with vehicle (grey), 5 mg/kg PF-9363 (orange), 0.033% SNDX-5613 (green) or combination (combo, 5 mg/kg PF-9363 and 0.033% SNDX-5613, purple).

**f, g, h** Statistics were calculated using a two-tailed t-test comparing means of each treatment group against each other (****p < 0.0001, ***p < 0.001, **p < 0.01, *p < 0.05, ns = p > 0.05)

**Supplemental Figure 6**

**a**, Experimental overview of *in vivo* single cell lineage tracing experiment using SPLINTR. Molm-13 cells were transplanted into NSG recipient mice (n=5 mice / group).

**b**, Kaplan-Meier survival curve of NSG mice transplanted with Molm-13 cells and treated with vehicle (grey), 5 mg/kg PF-9363 (orange), 10 mg/kg SNDX-5613 (SNDX, green) or combination (5 mg/kg PF-9363 and 10 mg/kg SNDX-5613, purple). Dosing commenced 4 days after transplantation and continued for the duration of the experiment. Log-rank (Mantel-Cox) test comparing survival of mice in the vehicle and combination treatment groups.

**c**, Boxplot of Shannon diversity index of SPLINTR barcodes in the baseline sample and across each treatment group.

**d**, Volcano plot of differential barcode abundance of clones in spleen of mice treated with combination versus vehicle bone marrow samples. Adjusted p-value < 0.05 and an absolute log_2_ fold change of 1.

**e**, Heatmaps of differential barcode abundance in the combination versus vehicle groups across the bone marrow (right) and spleen (left).

**f**, UMAP showing Louvain clusters across scRNA-seq experiment.

**g**, UMAP of vehicle and combination treatment groups annotated with responsive clones.

**h**, GSEA of MSigDB hallmark pathways enriched in cells from the combination treatment versus cells from the vehicle treatment. Positive scores indicate enrichment in the combination group. Negative scores indicate enrichment in the vehicle group.

**i**, GSEA plots of hallmark Oxidative Phosphorylation, E2F targets, MYC targets V2 and MYC targets V1 gene sets in the combination group versus the vehicle group. Negative scores indicate enrichment in the vehicle group.
