## Supplementary figures and images for "Catalytic inhibition of KAT6/KAT7 enhances the efficacy and overcomes primary and acquired resistance to Menin inhibitors in MLL leukaemia"

### Figure_S1

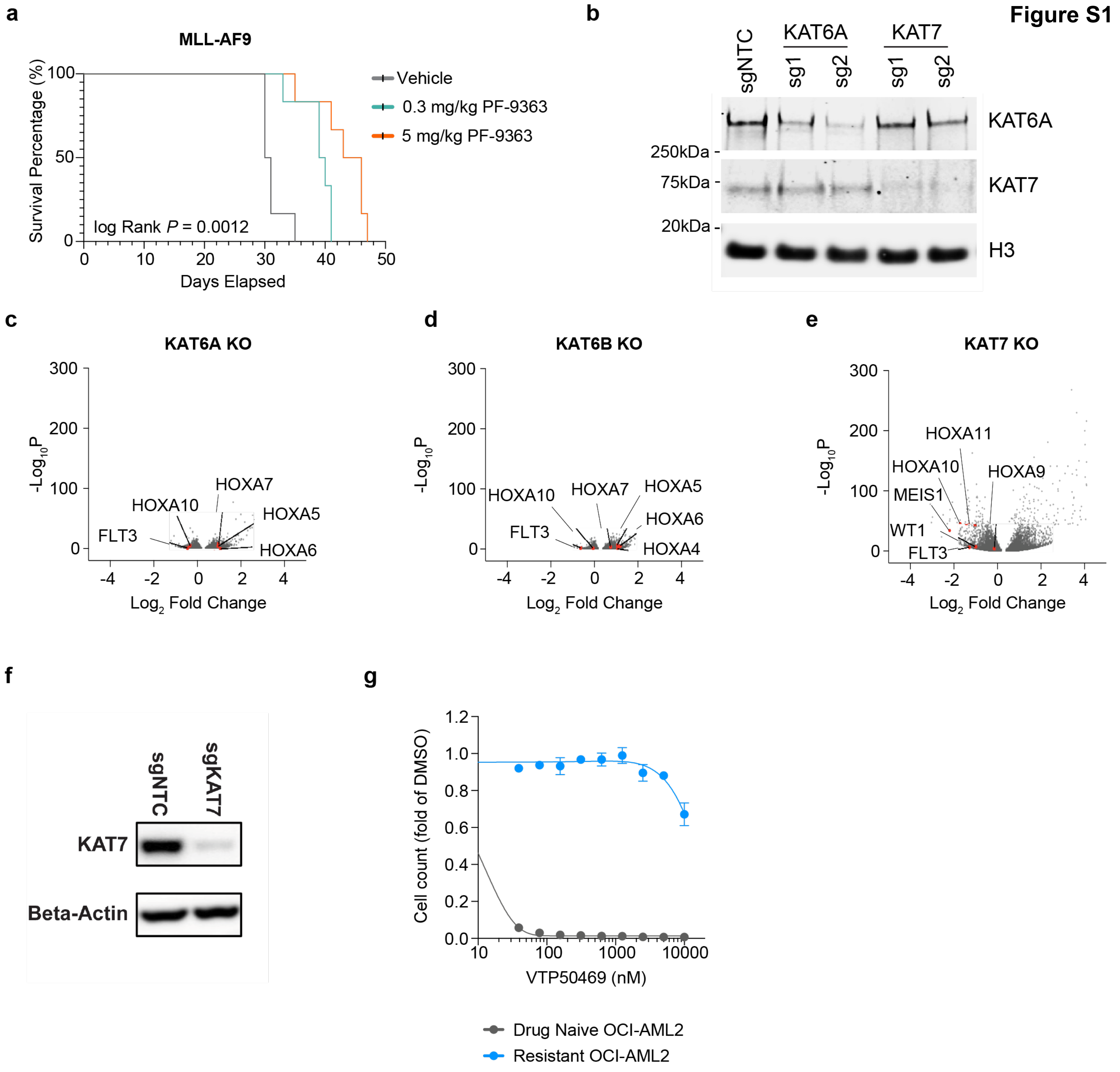

### Figure_S3

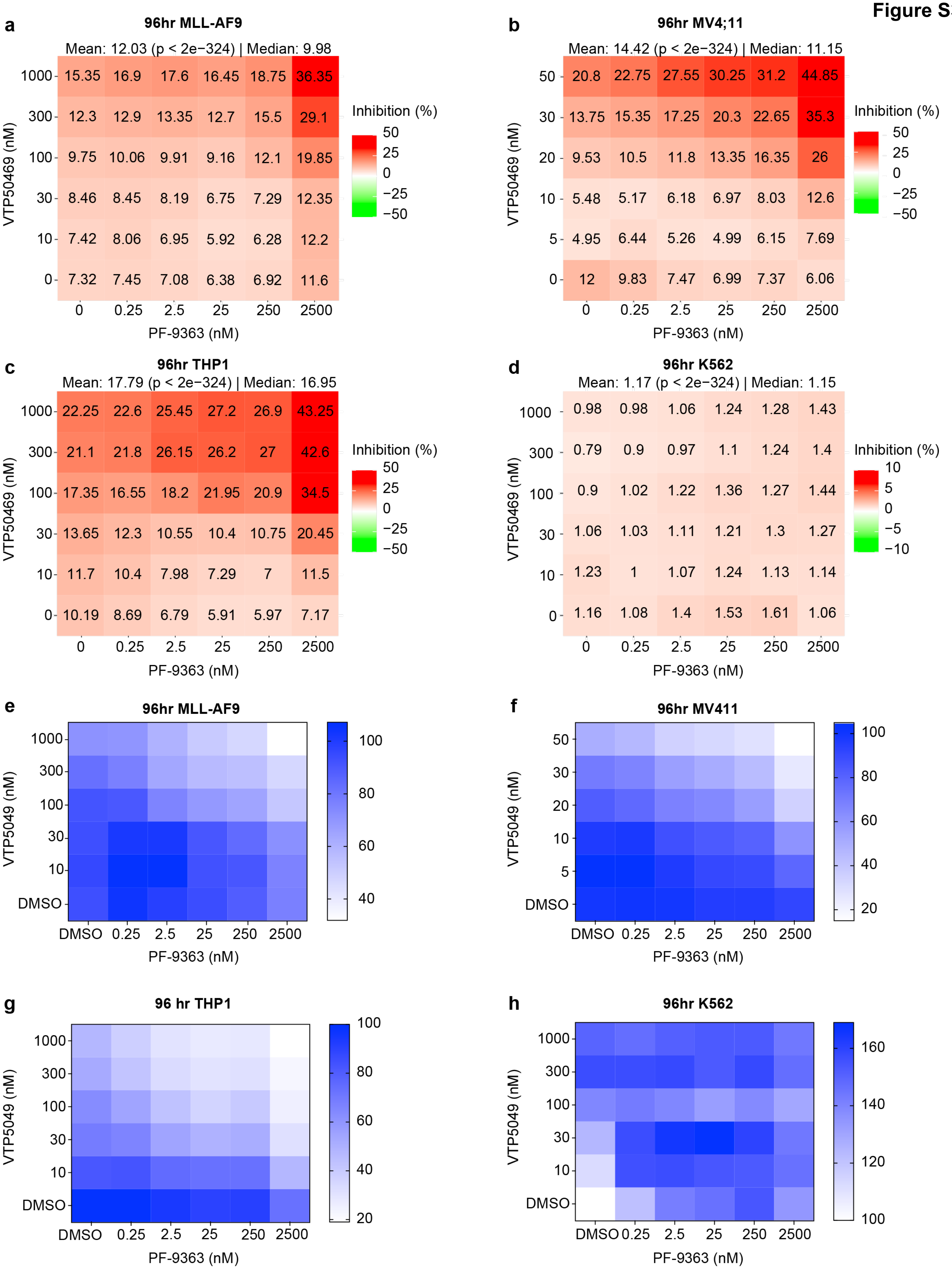

### Figure_S4

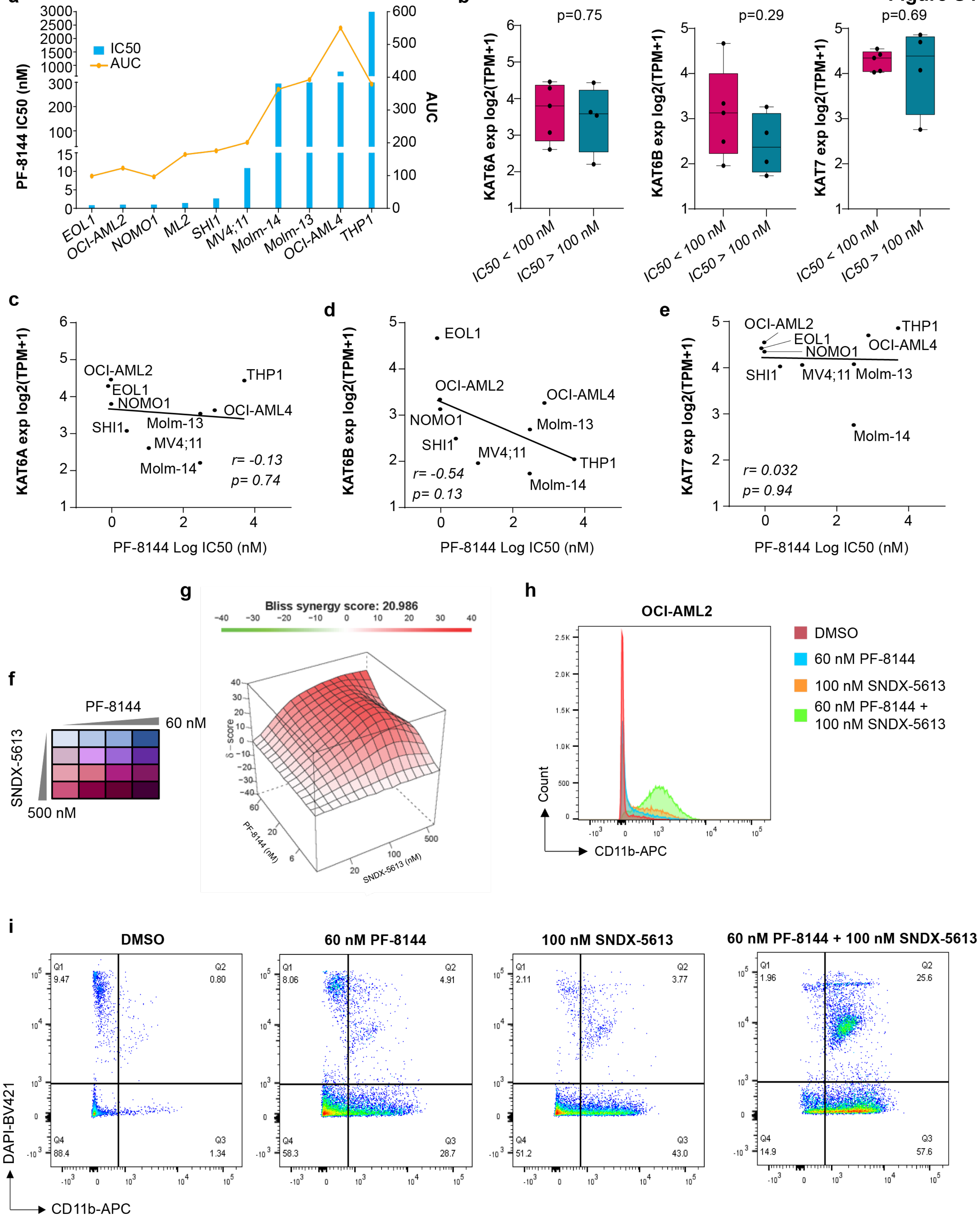

### Figure_S5

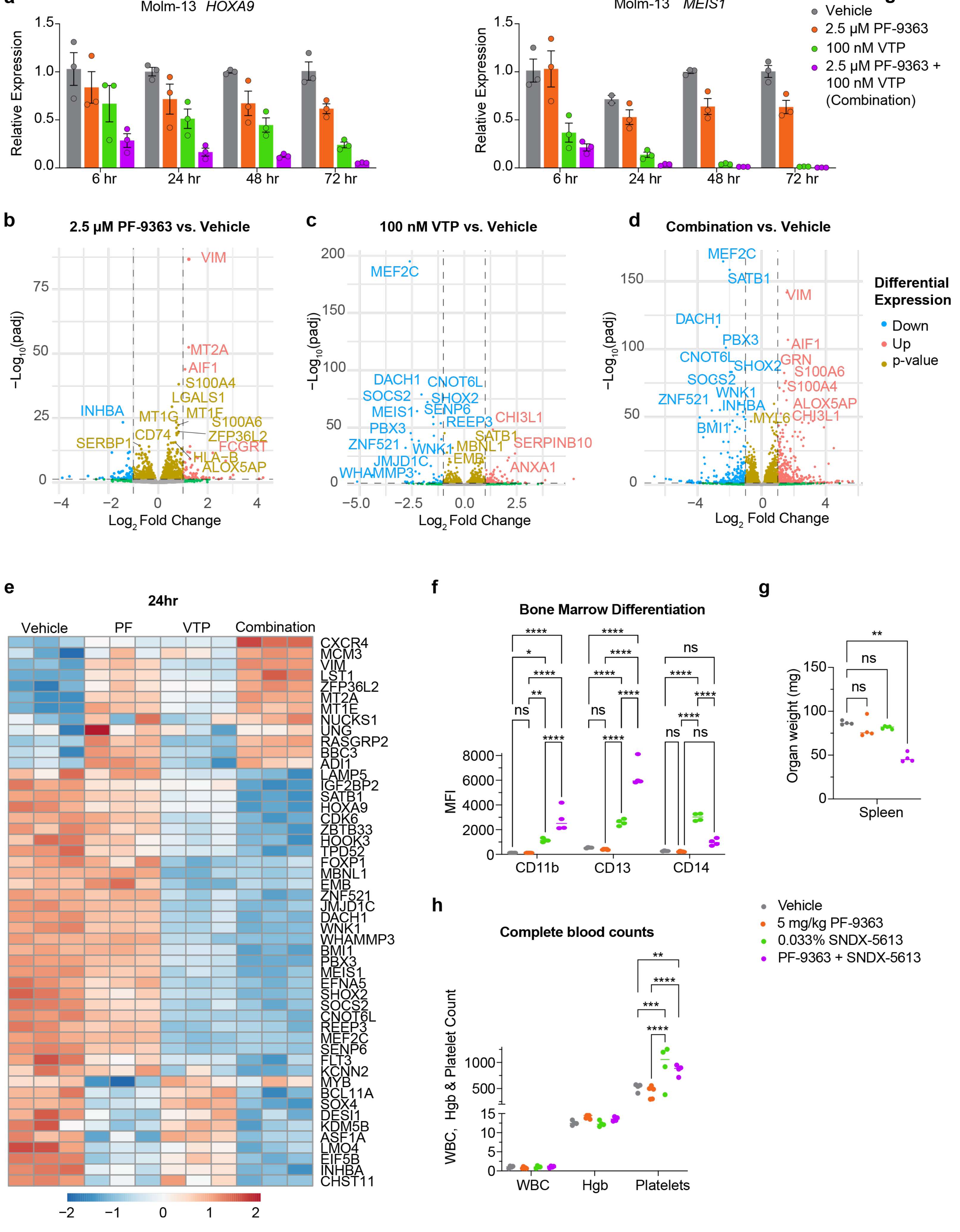

### Figure_S6

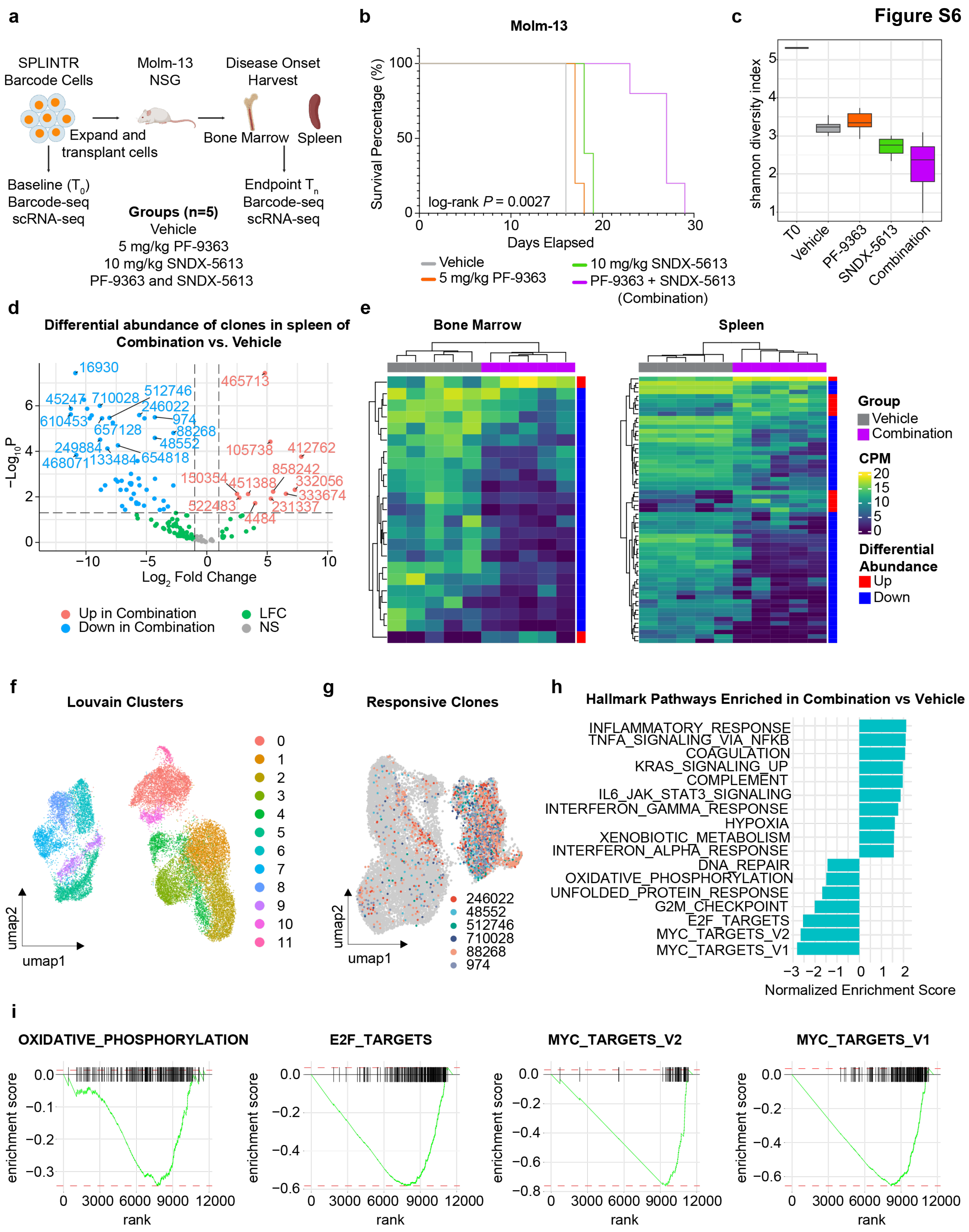
