## Supplementary material for "Catalytic inhibition of KAT6/KAT7 enhances the efficacy and overcomes primary and acquired resistance to Menin inhibitors in MLL leukaemia": Figure_S2

**a**

| Gene | Type | Protein | OCI-AML2-parental (VAF) | OCI-AML2-resistant-1 (VAF) | OCI-AML2-resistant-2 (VAF) |
|---|---|---|---|---|---|
| SPEN | DEL | p.T2892Ifs*2 | 0.45 | 0.48 | 0.47 |
| SPEN | SNP | p.K1600* | n.d. | 0.49 | 0.49 |
| SETDB1 | SNP | p.P5S | 0.52 | 0.49 | 0.48 |
| SESN3 | SNP | p.R457Q | 0.45 | 0.46 | 0.46 |
| KMT2A | SNP | p.K1751* | 0.49 | 0.47 | 0.52 |
| SMARCD1 | SNP | p.A5V | 0.44 | 0.46 | 0.51 |
| ATXN2 | SNP | p.P64A | 0.24 | n.d. | n.d. |
| EP400 | SNP | p.P1659L | 0.52 | 0.51 | 0.48 |
| FLT3 | SNP | p.A680V | 0.48 | 0.48 | 0.53 |
| FOXA1 | SNP | p.G378S | 0.49 | 0.48 | 0.51 |
| FOXA1 | SNP | p.L148V | 0.48 | 0.50 | 0.47 |
| ARID4A | SNP | p.N724S | 0.50 | 0.49 | 0.58 |
| KNSTRN | DEL | p.L210Rfs*21 | 0.50 | 0.51 | 0.47 |
| CD276 | SNP | p.G508R | 0.46 | 0.46 | 0.44 |
| CREBBP | SNP | p.R1446C | n.d. | 0.02 | n.d. |
| ZFHX3 | DEL | p.R3381_Q3388del | 0.48 | 0.49 | 0.42 |
| PLCG2 | SNP | p.E165K | 0.48 | 0.45 | 0.45 |
| FANCA | SNP | p.E1451K | 0.47 | 0.49 | 0.53 |
| TP53 | SNP | p.Y205S | 0.13 | 1.00 | 1.00 |
| FLCN | SNP | p.N184K | 0.48 | 0.47 | 0.50 |
| TYK2 | SNP | p.A53T | 0.50 | 0.46 | 0.45 |
| KMT2B | SNP | p.P679L | 0.09 | 0.49 | 0.50 |
| DNMT3A | SNP | p.R635W | 1.00 | 1.00 | 1.00 |
| GNAS | SNP | p.M162V | 0.45 | 0.45 | 0.49 |
| FAT1 | SNP | p.V295M | 0.53 | 0.44 | 0.50 |
| FAT1 | SNP | p.V129L | 0.49 | 0.50 | 0.51 |
| E2F3 | SNP | p.A376T | 0.31 | n.d. | n.d. |
| ROS1 | SNP | p.A1919D | 0.08 | n.d. | n.d. |
| PMS2 | SNP | p.M622I | 0.53 | 0.48 | 0.49 |
| HGF | SNP | p.A46V | 0.46 | 0.49 | 0.43 |
| VAV2 | SNP | p.I666M | 1.00 | 1.00 | 1.00 |
| ZRSR2 | INS | p.S447_R448dup | 0.77 | 0.73 | 0.75 |

**b**

| Gene | Type | Protein | PDX049D - parental (VAF) | PDX049D-resistant-1 (VAF) | PDX049D-resistant_2 (VAF) |
|---|---|---|---|---|---|
| CREBBP | INS | Q1079Sfs*8 | 0.476102941 | 0.462025316 | 0.49321267 |
| SMAD3 | SNP | I290= | 0.488069414 | 0.46969697 | 0.490196078 |
| TET2 | SNP | Y867H | 0.528541226 | 0.50997151 | 0.522357724 |
| VAV2 | SNP | I818M | 0.495918367 | 0.482837529 | 0.505725191 |
| TET2 | SNP | P1723S | 0.489296636 | 0.529166667 | 0.485250737 |
| ATM | SNP | P1054R | 1 | 1 | 1 |
| DIS3 | SNP | D438N | 0.474885845 | 0.444126074 | 0.46559633 |
| CREBBP | SNP | R714H | 0.541666667 | 0.450757576 | 0.44972067 |
| HLA-A | SNP | V189E | n.d. | 0.0625 | 0.067226891 |
| CSF3R | SNP | E808K | 0.494609164 | 0.525203252 | 0.50511509 |
| RAD51B | SNP | K243R | 0.468671679 | 0.514754098 | 0.495073892 |
| SPEN | SNP | D2007E | 0.4825 | 0.500777605 | 0.464922711 |
| PMS2 | SNP | T597S | 0.501838235 | 0.501187648 | 0.468864469 |
| MDC1 | SNP | E1536A | 0.485743381 | 0.468057366 | 0.470359572 |
| PIK3R2 | SNP | A727T | 0.52 | n.d. | 0.537037037 |
| SETD2 | SNP | T1077A | 0.488773748 | 0.463829787 | 0.493902439 |
| NSD1 | SNP | M2261T | 0.461424332 | 0.503496503 | 0.497576737 |
| CD58 | SNP | I237V | 0.489082969 | 0.484076433 | 0.412037037 |
| HLA-A | SNP | H216Y | 0.405797101 | 0.413407821 | 0.471153846 |
| HLA-A | SNP | F133L | 0.668989547 | 0.722222222 | 0.679054054 |
| CSF1R | SNP | E557K | 0.498888889 | 0.543189369 | 0.473132372 |
| EPHB1 | SNP | R367H | 0.519924099 | 0.488431877 | 0.486538462 |
| IRS2 | SNP | H1147Q | 0.483979764 | 0.481481481 | 0.488599349 |
| PRDM1 | SNP | R302P | 0.484149856 | 0.54863388 | 0.488829312 |
| ROS1 | SNP | V849F | 0.442260442 | 0.473239437 | 0.478589421 |
| ZFHX3 | DEL | Q3203_Q3204del | 0.376 | 0.380410023 | 0.372093023 |
| NSD1 | SNP | M2250I | 0.458823529 | 0.488425926 | 0.507836991 |
| NSD1 | SNP | A1036P | 0.490804598 | 0.521557719 | 0.494773519 |
| PPP4R2 | SNP | S282C | 0.492307692 | 0.5 | 0.489795918 |
| NSD1 | SNP | A691T | 0.495251018 | 0.535315985 | 0.47277937 |
| HLA-A | SNP | Q67R | 0.453674121 | 0.445783133 | 0.442073171 |
| APC | SNP | G2502S | 0.488304094 | 0.51048951 | 0.525830258 |
| PRKCI | SNP | R327= | 0.517899761 | 0.491525424 | 0.509054326 |
| HLA-C | SNP | A348V | 0.374558304 | 0.313559322 | 0.409090909 |
| FOXA1 | SNP | L148V | 0.476 | 0.517699115 | 0.477366255 |
| RTEL1 | SNP | T144I | 0.518796992 | 0.517316017 | 0.515769944 |
| KSR2 | SNP | R554Q | 0.516569201 | 0.493403694 | 0.512333966 |
| RECQL4 | SNP | P879H | 0.465240642 | 0.4743083 | 0.497326203 |
| PIK3CD | SNP | T456A | 0.537815126 | 0.479238754 | 0.475106686 |
| MEN1 | SNP | T349M | n.d. | 0.514440433 | n.d. |
| MEN1 | SNP | G331D | n.d. | n.d. | 0.4765625 |
